# SARS-CoV-2 entry route impacts a range of downstream viral and cellular processes

**DOI:** 10.1101/2022.08.05.502936

**Authors:** Bingqian Qu, Csaba Miskey, André Gömer, Dylan Potmus, Maximillian Nocke, Robin D.V. Kleinert, Tabitha K. Itotia, Lara Valder, Janice Brückmann, Sebastian Höck, Florian D. Hastert, Christine von Rhein, Christoph Schürmann, Aileen Ebenig, Marek Widera, Sandra Ciesek, Stephanie Pfaender, Zoltán Ivics, Barbara S. Schnierle, Alexander W. Tarr, Christine Goffinet, Michael D. Mühlebach, Daniel Todt, Richard J.P. Brown

## Abstract

SARS-CoV-2 entry is promoted by both cell-surface TMPRSS2 and endolysosomal cathepsins. To investigate the impact of differentially routed virions on host and viral processes, lung epithelial cells expressing distinct combinations of entry factors were infected with authentic viruses. Entry route determined early rates of viral replication and transcription, egress and inhibitor sensitivity, with differences observed between virus strains. Transcriptional profiling revealed that induction of innate immunity was correlated to viral genome and transcript abundance in infected cells. Surface entry triggered early activation of antiviral responses, reducing cumulative virion production, while endolysosomal entry delayed antiviral responses and prolonged virus shedding due to extended cell viability. The likely molecular footprints of escape from antiviral effector targeting were also recorded in viral genomes and correlated with entry route-dependent immune status of cells. TMPRSS2 orthologues from diverse mammals, but not zebra fish, facilitated infection enhancement, which was more pronounced for ancestral strains. Leveraging RNA-seq and scRNA-seq datasets from SARS-CoV-2 infected hamsters, we validate aspects of our model *in vivo*. In summary, we demonstrate that distinct cellular and viral processes are linked to viral entry route, collectively modulating virus shedding, cell-death rates and viral genome evolution.

## Introduction

The severe acute respiratory syndrome coronavirus 2 (SARS-CoV-2) is the etiological agent of the current COVID-19 pandemic. This originally zoonotic virus is readily transmissible between humans via the respiratory route. Indeed, despite extensive containment efforts since its initial spillover from an unidentified animal source to humans in late 2019 (***Zhou et al., 2020***), SARS-CoV-2 has spread globally. Infection results in a broad variety of clinical outcomes, ranging from asymptomatic to severe pneumonia and associated immune dysregulation that can lead to fatal multi-systemic failure (***García, 2020***). This range of disease severity observed between individuals is due to differential elicitation of host immune and inflammatory responses, and higher death rates are observed in the elderly reflecting immunosenescence (***Camell et al., 2021***). The continuous emergence of transmission enhancing mutations, mainly in the spike glycoprotein (S), has given rise to multiple variants of concern (VOC), possibly in response to the altered immune profiles in humans induced by natural infection and vaccination (***Harvey et al., 2021***). Omicron (B.1.1.529) has replaced Delta (B. 1.617.2) as the dominant circulating variant worldwide, with subsequent replacement of the original BA.1 sub-lineage with the more transmissible BA.2, BA.4 and BA.5 sub-lineages. Together B.1.1.529 lineage viruses and their decedents exhibit significant immune escape from vaccine-induced neutralizing antibodies (***Wilhelm et al., 2022***) and milder pathogenicity owing to an altered cell tropism (***Fan et al., 2022***).

Despite the diversity of S proteins between variants, a critical point of the SARS-CoV-2 lifecycle is the entry step, which appears to be conserved. During virion assembly in infected cells, furin processing in the trans-golgi network results in S1-S2 cleavage, exposing the C-terminus of S1 for further proteolytic processing (***Daly et al., 2020*; *Kielian, 2020***). SARS-CoV-2 infection of new cells initiates via cell attachment which involves C-type lectin receptors (e.g. L-SIGN, DC-SIGN and SIGLEC1) (***Lempp et al., 2021***). The virus attaches to cellular membranes and direct binding of the S protein to the primary receptor, angiotensin converting enzyme 2 (ACE2), precedes membrane fusion (***Hoffmann et al., 2020*; *Zhou et al., 2020***). After ACE2 engagement, which induces conformational changes in the S protein, there are two distinct cellular locations at which virus-host membrane fusion occurs (***Jackson et al., 2022***). S protein cleavage at the S2’ site exposes the fusion peptide facilitating fusion of viral-host membranes, creating a pore through which the viral genome is released into the cellular cytoplasm. In the presence of human transmembrane protease serine 2 (hTMPRSS2) cleavage of S2’ occurs at the cellular surface. Under these conditions, the virus enters the cell within 10 mins in a pH independent manner (***Koch et al., 2021***). On the other hand, if the cell expresses insufficient amounts of TMPRSS2, the entire virus-hACE2 complex is internalized via clathrin-mediated endocytosis into endolysosomes, where S cleavage is mediated by cathepsin L (***Zhao et al., 2021***). This step takes 40-60 mins post initial attachment (***Bayati et al., 2021*; *Koch et al., 2021***). Therefore, cell-surface TMPRSS2 expression levels determine which pathway the virus utilizes to enter permissive ACE2 expressing cells.

TMPRSS2 was first identified as a host protease responsible for influenza virus hemagglutinin cleavage. It belongs to the family of type II transmembrane serine proteases and contains an N-terminal cytoplasmic tail, a transmembrane domain (TM), a low-density lipoprotein receptor class A domain (LDLa), a scavenger receptor cysteine-rich domain (SRCR), and a C-terminal trypsin-like serine protease domain consisting of a catalytic triad: histidine 296, aspartic acid 345 and serine 441 (HDS) (**Fig. 1A**) (***Böttcher-Friebertshäuser et al., 2010***). The TMPRSS2 protein can be auto-processed or processed by other proteases at the R255-I256 junction. TMPRSS2 is not only expressed in airway epithelia, but also in small intestinal enterocytes. The presence of TMPRSS2 in gut activates spike fusogenic activity and facilitates viral entry into intestinal cells (***Zang et al., 2020***). TMPRSS2 also promotes the entry process of a broad number of coronaviruses, including HCoV-229E, SARS-CoV and MERS-CoV in a cathepsin L-independent manner (***Bertram et al., 2013*; *Gierer et al., 2013*; *Shulla et al., 2011***). Therefore, TMPRSS2 represents a promising host target for antiviral interventions and therapeutic targeting of TMPRSS2 by small molecule inhibitors has been reported. Using a SARS-CoV-2 pseudoparticle assay, camostat mesylate was identified as a potential entry inhibitor, with *in vitro* EC_50_ and EC_90_ values of approximately 1 μM and of 2-5 μM, respectively (***Hoffmann et al., 2020***). Camostat mesylate, its metabolite 4-(4-guanidinobenzoyloxy) phenylacetic acid, and a related protease inhibitor, nafamostat mesylate, also exhibited antiviral activity in human lung slices *ex vivo* (***Hoffmann et al., 2021***). Mechanistically, camostat mesylate is likely inserted into the pocket within TMPRSS2 responsible for the binding of S and TMPRSS2, and covalently binds to residues on TMPRSS2 and thereby blocks its catalytic activity (***Sun et al., 2021***). Small-molecule inhibitor lead MM3122 (ketobenzothiazole) is more potent than either camostat or nafamostat, with an EC_50_ of 430 pM for inhibition of pseudovirus entry and 74 nM for inhibition of authentic-virus entry to Calu-3 cells (***Mahoney et al., 2021***). Recently, small molecule compound N-0385 was identified, which has a nanomolar EC_50_ value against SARS-CoV-2 infection in human lung cells and colonoids. Notably, this compound inhibited entry of variants of concern B.1.1.7, B.1.351, P.1 and B.1.617.2 *in vitro* and reduced viral loads in infected K18 human ACE2-transgenic mice (***Shapira et al., 2022***). Collectively, these pre-clinical findings confirm TMPRSS2 represents a viable therapeutic target in the host.

**Figure 1:**
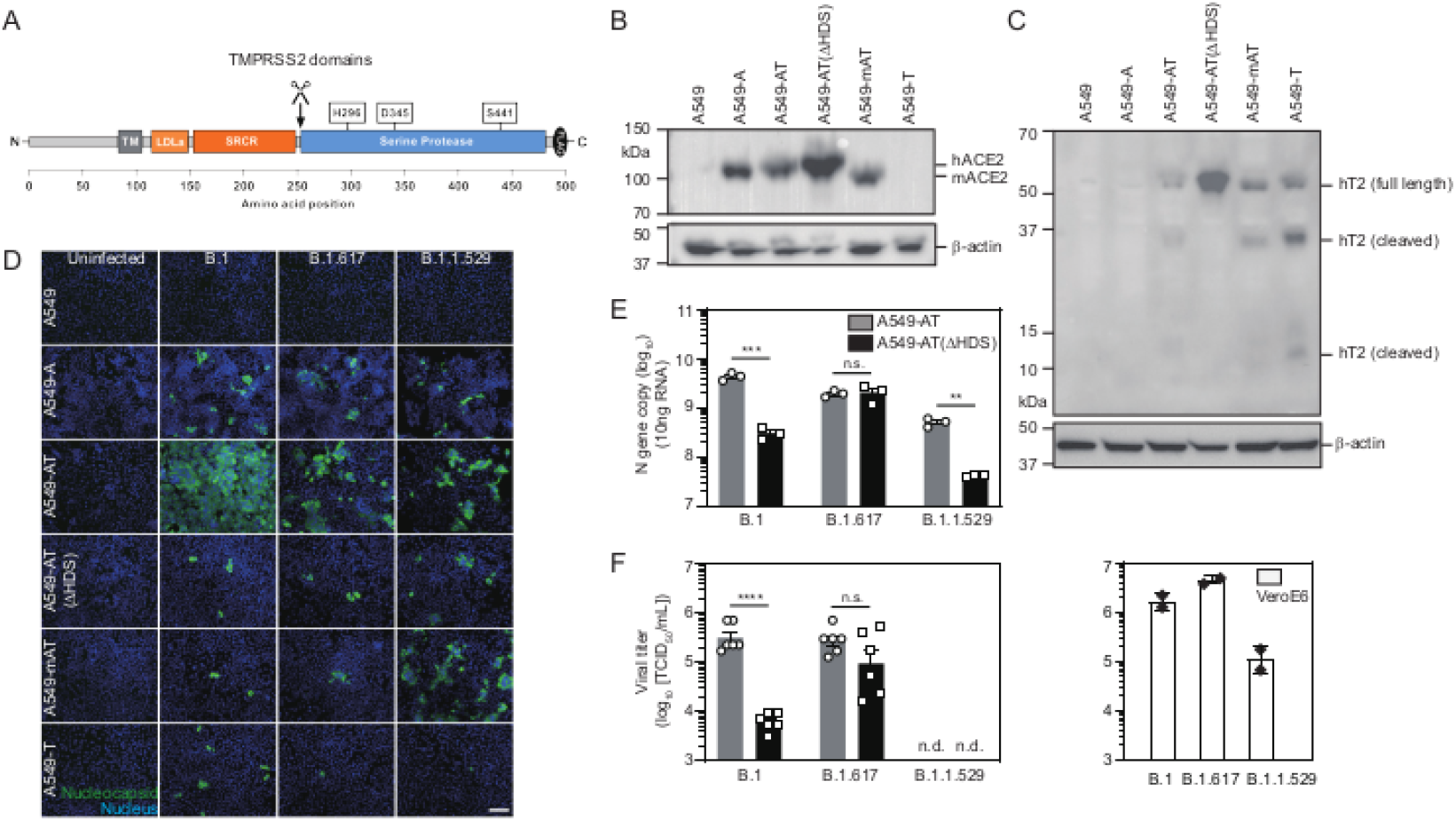
Human TMPRSS2 expression increases SARS-CoV-2 replication and virion egress in a strain dependent manner. **(A**) Cartoon of human TMPRSS2 domains. Location of catalytic triad residues and auto-cleavage site (scissors) are highlighted above. (**B**) Human or murine ACE2 protein expression in the indicated cell lines, determined by Western blot analysis. (**C**) Human TMPRSS2 protein expression levels in the indicated cell lines, determined by Western blot analysis. Bands representing full-length protein and two fragments formed by auto-cleavage are highlighted. (**D**) Susceptibility of indicated cell lines to B.1, B.1.617 and B.1.1.529 infection (MOI=0.01, 24 hpi). Immunofluorescence staining of SARS-CoV-2 nucleocapsid protein (green) or cellular nuclei (blue). Scale bar: 200 μM. (**E**) Human TMPRSS2 enhances replication and transcription of B.1 and B.1.1.529 but not B.1.617. Intracellular *N* copy numbers per 10ng total RNA (MOI=0.01, 24 hpi, n=3 ± SEM). (**F**) Human TMPRSS2 increases the secretion rates in a strain dependent manner. Left panel: Viral titers in the medium of A549-AT and A549-AT(ΔHDS) cells at 24 hpi were determined using TCID_50_ (n=6 ± SEM). Right panel: Virion secretion of the indicated strains from VeroE6 cells (MOI=0.01, 24 hpi, n=2 ± SEM). ****P<0.0001, ***P<0.001, **P<0.01, n.s.: no significance. n.d.: not detected.

Highly pathogenic viruses have traditionally been cultured using the VeroE6 primate kidney endothelial cell-line due to the high titers produced. This is partially due to defective innate immunity, which includes an absence of baseline interferon stimulated gene (ISG) expression and an inability to produce interferon (IFN) upon infection (***Felgenhauer et al., 2020***). To investigate a link between SARS-CoV-2 entry route and physiologically relevant host and viral processes downstream, we utilized A549 cells in this study as they possess a number of favorable properties. These include: (1) a tissue and species origin that is biologically relevant (human lung epithelial cells), (2) an absence of endogenous ACE2 receptor and TMPRSS2 accessory protease expression, allowing controlled ectopic supplementation, and (3) maintenance of a functional IFN-system (***Widera et al., 2021***), including the ability to produce IFN and subsequent induction of downstream effectors. We used these cells to investigate a number of questions including whether ancestral and VOC SARS-CoV-2 strains differ in their TMPRSS2 dependence and susceptibility to inhibitors, whether TMPRSS2- or cathepsin L-routed virus differentially activate host innate immune responses, whether entry route impacts virion production rates and lifespan of infected cells, whether entry route impacts mutation frequencies in viral populations, and finally, whether SARS-CoV-2 entry enhancement is a conserved property of TMPRSS2 orthologues from diverse species.

## Results

### Characterization of ACE2- and TMPRSS2-expressing A549 cell-lines

We engineered lentiviral transgene delivery plasmids encoding either the human or murine ACE2 proteins, in addition to lentiviral transgene delivery plasmids encoding either a functional C-terminally flag-tagged human TMPRSS2 protein (**Fig. 1A**) (***Böttcher-Friebertshäuser et al., 2010***) or a catalytically inactive version (ΔHDS). Plasmids were used to generate a panel of ectopically expressing cell-lines, and Western blot analysis demonstrated human or murine ACE2 expression in A549-A, A549-AT, A549-AT(ΔHDS) and A549-mAT cells (**Fig. 1B**). Human TMPRSS2 was expressed and auto-cleavage of human TMPRSS2 generated two fragments (32 kD and 12 kD) in A549-AT and A549-T cells. Accumulated full-length TMPRSS2 was observed in A549-AT(ΔHDS) cells, in which serine protease activity and therefore auto-cleavage, was abolished (**Fig. 1C**). These analyses confirm no endogenous protein expression for ACE2 or TMPRSS2 in parental A549 cells and verify ectopic expression in engineered cell-lines.

### TMPRSS2 expression increases susceptibility to B.1 and B.1.1.529 viruses

Next, we evaluated the susceptibility of parental or engineered cell-lines to infection with ancestral virus (B.1) and VOCs (B.1.617 and B.1.1.529). All three strains replicated at a comparable level in A549-A cells, as determined by immunofluorescence staining of nucleocapsid protein (**Fig. 1D**). Replication of B.1 was markedly enhanced in A549-AT cells, and modestly increased for VOCs. Enhancement of all three viruses was reduced to levels comparable to, or lower than, A549-A cells when TMPRSS2 was catalytically inactive (**Fig. 1D**). Remarkably, B.1.1.529 nucleocapsid protein accumulation was similar in TMPRSS2 cells overexpressing either human or murine ACE2, which was not the case for B.1 and B.1.617 variants, implying an expansion of host range to utilize murine ACE2 for B.1.1.529. Intracellular replication and transcription of B.1 and B.1.1.529 were significantly enhanced in A549-AT compared to A549-AT(ΔHDS) cells, but this was not observed for B.1.617 (**Fig. 1E**). The picture was again distinct for virion secretion rates, which were significantly enhanced in A549-AT for B.1, modestly enhanced in A549-AT for B.1.617 and undetectable in supernatants from both in A549-A and A549-AT for B.1.1.529. (**Fig. 1F,** left panel). In contrast, robust B.1.1.529 egress in VeroE6 cells was confirmed, albeit at lower rates than B.1 and B.1.617 (**Fig. 1F,** right panel). These data confirm significant differences in TMPRSS2 dependence and lung epithelial cell tropism between SARS-CoV-2 strains.

### TMPRSS2 expression modulates SARS-CoV-2 entry inhibition

To explore susceptibility to entry inhibition in lung epithelial cells which allow distinct entry pathways, we performed authentic virus infections in the presence of camostat mesylate, a TMPRSS2 inhibitor, and E-64d, a cathepsin B/L inhibitor. Vacuolar H^+^ ATPase inhibitor bafilomycin A1 was used as positive control. (**Fig. 2A-2D and Supplementary Fig S1**). Camostat mesylate modestly inhibited B.1 infection in A549-AT cells (1 log reduction in virion production), but not A549-A cells. B.1.617 infection of either cell type was not inhibited. In contrast, E-64d significantly inhibited B.1 and B.1.617 infection of both cell-types. Of note, B.1 infection of A549-AT(ΔHDS) cells was further reduced under all conditions when compared to A549-A cells, implying functionally dead TMPRSS2 actively blocks ACE2 engagement by steric hindrance. This effect was not observed for B.1.617, highlighting potential differences in ACE2 engagement between strains. Additionally, B.1.617 was able overcome vacuolar H^+^ ATPase inhibition via TMPRSS2-mediated surface entry whereas B.1 was not. These data point to subtle differences in receptor engagement and TMPRSS2 dependence between strains which regulate sensitivity to inhibition.

**Fig 2:**
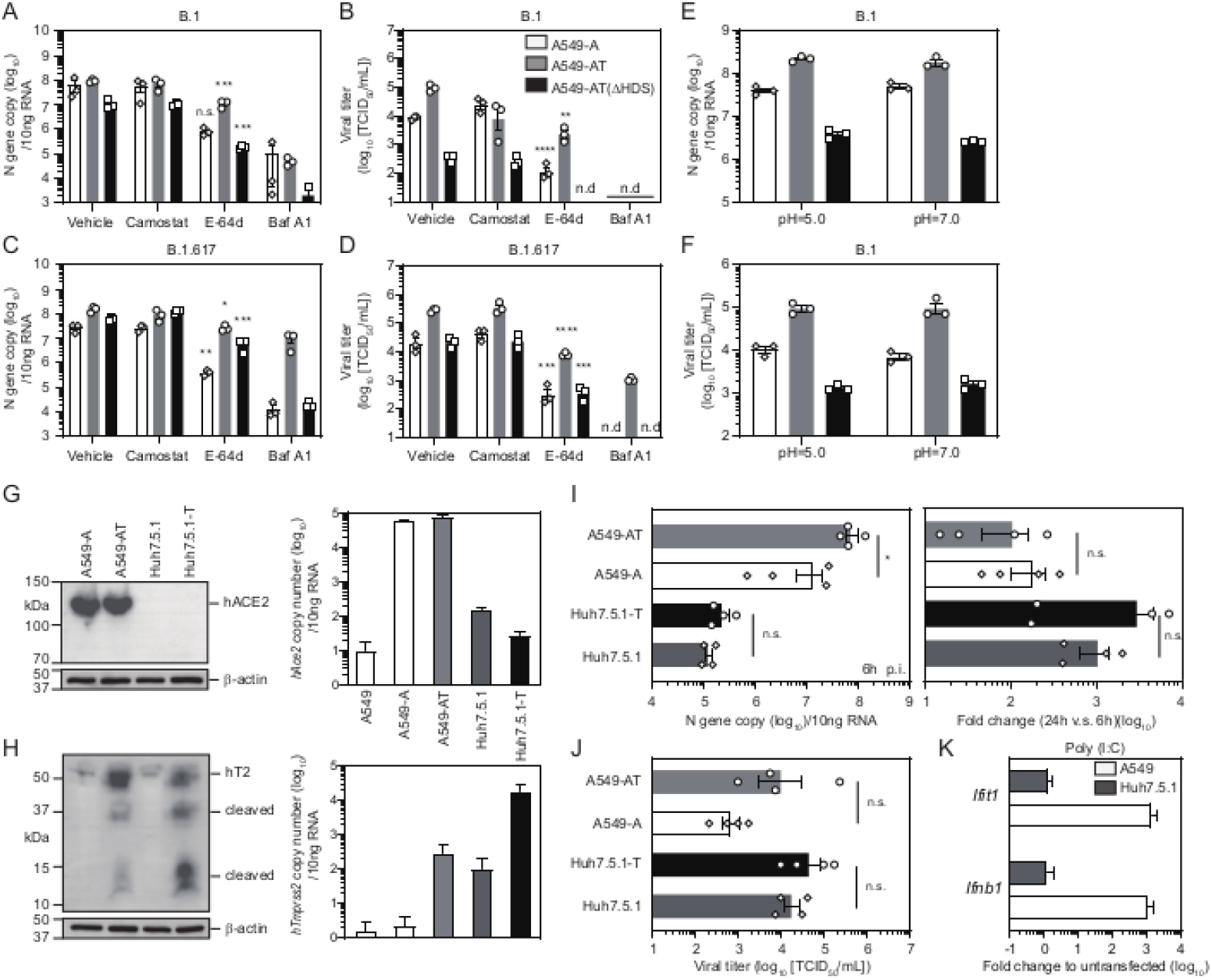
TMPRSS2 expression modulates sensitivity to pharmacological and innate immunity inhibition. (**A** and **B**) Inhibition of B.1 entry. (**A**) Intracellular *N* gene RNA copies and (**B**) virion secretion in the indicated cell-lines after infection in the presence of inhibitors (MOI=0.01, 24 hpi, n=3 ± SEM). Baf A1: bafilomycin A1. (**C** and **D**) Inhibition of B.1.617 entry. (**C**) Intracellular *N* gene RNA copies and (**D**) virion secretion in the indicated cell-lines after infection in the presence of inhibitors (MOI=0.01, 24 hpi, n=3 ± SEM). (**E** and **F**) Environmental pH does not mediate fusion peptide exposure. (**E**) Intracellular *N* gene RNA copies and (**F**) B.1 secretion in the indicated cell-lines after infection at the stated pH (MOI=0.01, 24 hpi, n=3 ± SEM). Endogenous and ectopic ACE2 expression in A549 and Huh-7.5.1 cells. (**G**) Human ACE2 protein (left) and RNA transcript levels (right) in the indicated cell-lines. (**H**) Human TMPRSS2 protein (left) and transcript levels (right) in the indicated cell-lines. (**I**) Innate immunity correlates with impaired viral replication rates. The indicated cell-lines were infected with B.1 at an MOI of 0.01. Left: intracellular *N* gene RNA at 6 hpi. Right: relative fold change in vRNA levels at 24 hpi compared to 6 hpi (n=4 ± SEM). (**J**) Virion secretion is enhanced in the absence of innate immunity. Virion secretion at 24 hpi in the cells from (**I**). (**K**) In contrast to A549 cells, Huh7.5.1 cells do not upregulate IFN or ISGs upon Poly(I:C) stimulation. *P<0.05, **P<0.01, ***P<0.001, ****P<0.0001, n.s.: no significance. n.d.: not detected.

Many virus families hijack clathrin-mediated endocytosis pathways to enter cells, with virus-host membrane fusion occurring in acidified endosomes due to pH-induced conformational changes which expose the fusion peptide. To investigate whether surface entry at the plasma membrane can be artificially induced in TMPRSS2-deficient cells, we performed infections under acidified conditions (**Fig. 2E-2F**). These data indicate that surface entry of SARS-CoV-2 cannot be induced by low pH and confirm that fusion peptide exposure is protease-mediated and not regulated by pH-induced conformational rearrangements.

### Innate immunity suppresses viral propagation

We observed that the human hepatoma cell-line Huh7.5.1 is susceptible to B.1 infection. As Huh7.5 cells and their derivatives are deficient in innate immune signaling (***Tegtmeyer et al., 2021***) we reasoned this cell-line would be a useful tool to compare entry routes in the absence of innate immune suppression. ACE2 and TMPRSS2 protein expression were not detected in Huh7.5.1 cells by Western blot analysis, although endogenous *ACE2* and *TMPRSS2* transcript levels were higher than in A549 cells, and ectopic TMPRSS2 expression significantly boosted both protein and RNA levels (**Fig. 2G and 2H**). After B.1 infection at 6 hpi, intracellular viral RNA levels in Huh7.5.1-T cells were ~300-fold less than A549-AT cells (**Fig. 2I Left**). However, vRNA levels at 24 hpi versus 6 hpi in Huh7.5.1-T were ~30-fold increased when compared to A549-AT cells, suggesting more robust viral replication due to a lack of innate immunity (**Fig. 2I Right and 2J**). Indeed, we confirmed parental Huh7.5.1 cells were unresponsive to Poly(I:C) stimulation while robust induction was observed in A549 cells (**Fig. 2K**). Despite higher initial uptake in A549-AT cells when compared A549-A cells, fold change in viral RNA levels in A549-A cells were modestly increased when compared to A549-AT (6 vs 24 hpi). This result hints that the extent of innate immune induction and concomitant suppression of viral replication in engineered A549 cells may be modulated by entry route. Together these data confirm SARS-CoV-2 can enter cells with low endogenous ACE2 expression and that intact innate immunity correlates with suppressed viral propagation.

### SARS-CoV-2 entry route determines downstream viral genome replication and transcription rates

SARS-CoV-2 genomic and subgenomic RNAs are flanked by a 5’ cap and a 3’ polyA tail (***Kim et al., 2020***). Therefore, RNA-sequencing (RNA-seq) of SARS-CoV-2 infected cells recovers both host and viral transcripts, allowing simultaneous investigation of viral replication and transcription rates, host responses to infection and patterns of viral substitutional evolution. Consequently, infections of A549-A and A549-AT cells were performed with our panel of viruses at low (0.01) or high (1) MOIs. At 24 hpi, differential virion production rates were observed in A549-A and A549-AT cells, which differed in magnitude between strains (**Fig. 3A** and **1F**). As previously described, differences in virus secretion rates between A549-A and A549-AT cells at low MOI were most pronounced for strain B.1 (~17-fold), while B.1.617 exhibited lower TMPRSS2-dependence (~4.5-fold difference). At high MOI, virus secretion rates were demonstrably increased although differences between A549-A and A549-AT cells were less pronounced, with either ~3-fold (B.1) or no differences (B.1.617) observed. In contrast, no virion release was detected for B.1.1.529 infections, irrespective of TMPRSS2 expression or MOI.

**Figure 3.**
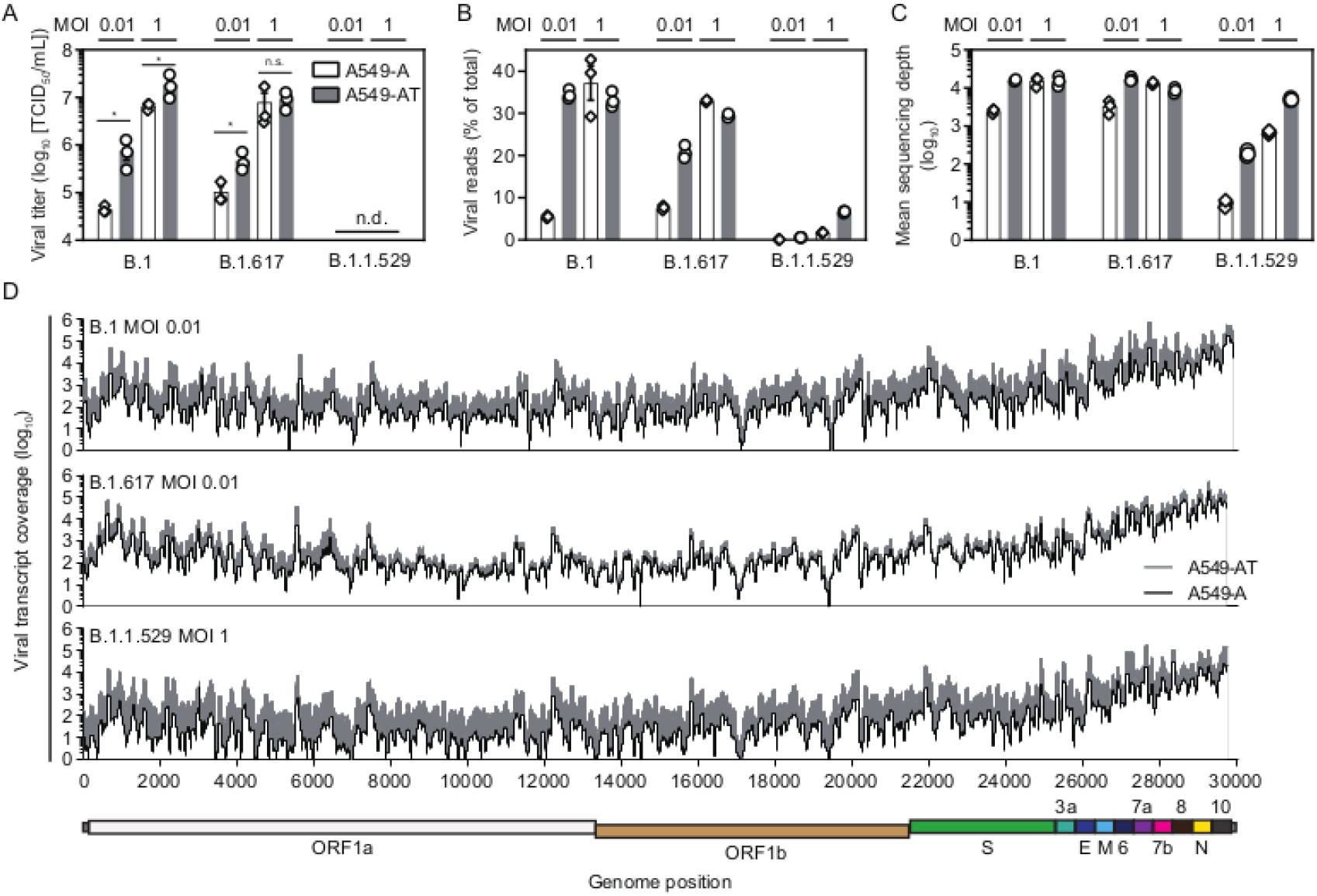
Differences in cell-resident viral transcript abundance after surface or endolysosomal entry. Infections of A549-A and A549-AT cells were performed with the indicated SARS-CoV-2 strains at low and high MOI, with supernatants and cellular RNAs harvested at 24 hpi. **(A)** Viral titres in harvested supernatants. **(B**) Number of virus-genome mapped reads as a percentage of total reads. **(C)** Mean SARS-CoV-2 genome/transcript sequencing depth. **(D)** Direct comparison of mean genome/transcript coverage (n=3) across the respective SARS-CoV-2 genomes in A549-A and A549-AT cells. Cartoon directly below represents the relative locations of genome encoded ORFs.

To determine the effect of viral entry route on downstream levels of viral replication and transcription, we performed RNA-seq on cellular RNAs at 24 hpi. Reads were mapped against the respective viral reference genome and the human *hg38* genome scaffold, and the ratio of viral reads to total cellular reads calculated (**Fig. 3B**). At low MOI, a 6.4-fold increase in B.1 transcripts was observed between A549-A and A549-AT cells (5.5% versus 34.5%), compared to a ~2.7-fold increase for B.1.617 (7.6% versus 20.8%). In contrast, at high MOI, minimal differences were observed between B.1 and B.1.617 transcript levels in A549-A and A549-AT cells, indicating viral entry and downstream replication rates have reached saturation point in this cell type. At high MOI, B.1 transcript levels were not elevated above low MOI levels in A549-AT cells, although a ~10% increase was observed for B.1.617 transcripts. Despite abundant ACE2 expression in both A549-A and A549-AT cells, B.1.1.529 transcript levels were drastically reduced when compared to B.1 and B.1.617. However, a ~6.2-fold increase was apparent between A549-A and A549-AT cells at low MOI (0.09% versus 0.58%) while a ~3.9-fold difference was observed at high MOI (1.72% versus 6.64%), indicating B.1.1.529 entry is also enhanced by TMPRSS2 expression.

Mean sequencing depth (**Fig. 3C**) and read coverage across viral genomes (**Fig. 3D**) highlights differences in early viral replication and transcription rates apparent between A549-A and A549-AT cells. Independent of transcript depth or infecting strain, these analyses also revealed highly conserved transcript profiles tiled across the SARS-CoV-2 genome, confirming conservation of ORF expression and genome replication kinetics between ancestral and VOC strains. Taken together, these data confirm TMPRSS2 expression enhances infection rates of SARS-CoV-2 B.1, B.1.617 and B.1.1.529 strains, with B.1.617 exhibiting the least dependency. When surface entry occurs, greater numbers of viral genomes are delivered to the cytosol to seed initial infection, resulting in a concomitant elevation of viral replication and transcription rates. These data also confirm authentic B.1.1.529 entry is modulated by TMPRSS2 expression and demonstrate both reduced entry rates and complete ablation of virion egress for this VOC in A549 lung epithelial cells.

### TMPRSS2-routed SARS-CoV-2 entry promotes early activation of cellular antiviral programs

We reasoned viral genome/transcript levels in cells would impact the magnitude of infection-induced changes in the cell-intrinsic transcriptional landscape. As the SARS-CoV-2 genome encodes IFN antagonists (***Schroeder et al., 2021***) we also included parental A549 cells transfected with the dsRNA mimic Poly(I:C) to confirm the cell-line’s ability to mount antiviral responses without inhibition. Expression of relevant entry factors and components of the IFN system in parental cells without infection was confirmed, in addition to the ability to produce IFN upon Poly(I:C) stimulation. (**Supplementary Fig S2**). Principal component analyses (PCA) of individual transcriptomes revealed tight clustering of biological replicates, with segregation of individual clusters dependent on infection status, presence or absence of TMPRSS2 expression, infecting strain and MOI (**Fig. 4A**). After host-transcript mapping to the human *hg38* genome scaffold, statistical analyses identified differentially expressed genes (DEGs) induced upon SARS-CoV-2 infection of A549-A and A549-AT cells for all strains, or Poly(I:C) transfection of A549 cells. These analyses identified differences in the magnitude of cellular response to infection, which was broadly correlated to viral transcript abundance (**Supplementary Fig S3**). To visualize differential patterns of DEG induction between experimental conditions, we zoomed in on selected subsets of genes which are induced upon infection to suppress viral replication and reduce systemic dissemination by direct and indirect means, in addition to viral entry factors. (**Fig. 4B**).

**Figure 4:**
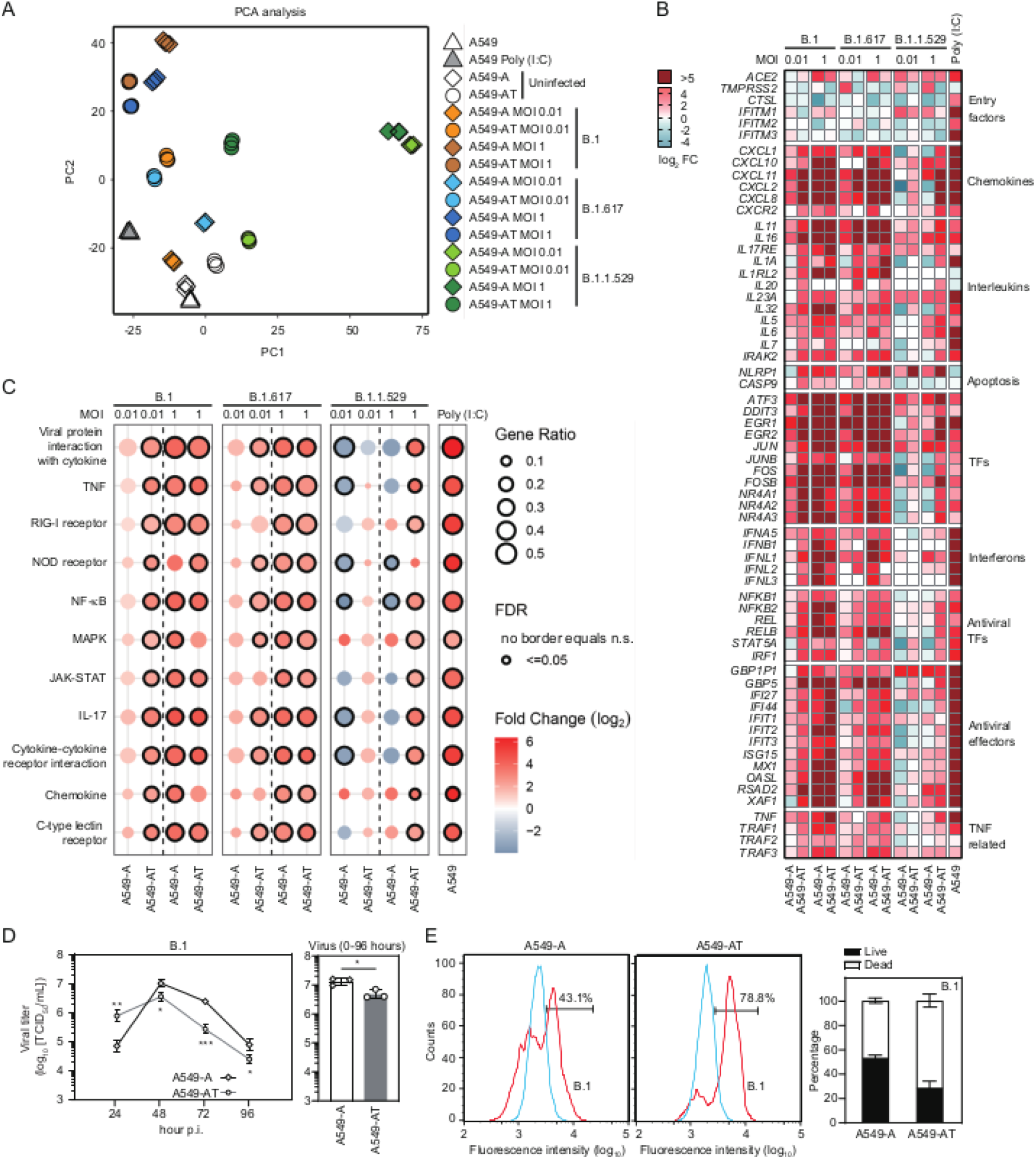
The magnitude of early antiviral responses is determined by SARS-CoV-2 entry route. **(A)** Principal component analysis (PCA) of the indicated transcriptomes. **(B)** Heat map visualizes fold change in mRNA expression (log_2_) for selected virus-inducible transcripts associated with different cellular processes after SARS-CoV-2 infection of A549-A and A549-AT cells. Fold change is relative to expression levels in the respective uninfected control cell-line. Data presented are mean values from n=3 biological replicates. **(C)** Dot-plot visualizes cell processes and signalling pathways targeted upon infection. FDR: false discovery rate. (**D**) Virus production rates in the indicated cell-lines (strain B.1, MOI 0.01). Left panel. Virion production kinetics 0-96 hpi. Right panel. Cumulative virus production. (**E**) Cell viability of the indicated cell-lines at 96 hpi. Left panels: example flow cytometric plots. Right panel. Percentages of live and dead cells. Blue: unstained, red: zombie dye staining.

Upon infection, expression of entry factors *ACE2* and *IFITM1* were generally upregulated while *TMPRSS2*, *CTSL*, *IFITM2* and *IFITM3* expression was broadly down-regulated. Induction of a subset of antiviral transcription factors (TFs), type I and III IFNs, and antiviral effector transcripts was directly correlated to cellular viral genome/transcript levels. At low MOI, minimal induction of these transcripts by B.1 or B.1.617 infection was observed in A549-A cells, while robust induction was apparent in A549-AT cells. A similar pattern of induction was observed for genes encoding chemokines, interleukins, apoptosis factors, tumour necrosis factor (TNF)-related transcripts and TFs including acute response genes and nuclear receptors (**Fig. 4B**). This effect was more pronounced for B.1 than B.1.617. The distinction between entry routes was lost at high MOI, where viral replication and transcription presumably reached saturation and robust induction was seen in both cell types. For B.1.1.529, robust induction was only observed at high MOI in A549-AT cells.

Infection-induced transcriptional dysregulation modulates a range of cell-intrinsic processes and canonical signalling pathways. We performed KEGG pathway analyses and confirmed that infection-induced changes in the cellular transcriptome mainly affected cellular processes associated with immunity and inflammation (**Fig. 4C**). Targeted cellular processes were generally conserved across strains, with significant activation correlated to viral genome/transcript abundance determined by entry route. Of note, pathways and processes activated upon B.1 and B.1.617 infection were differentially suppressed (A549-A) or activated (A549-AT) by B.1.1.529 infection (**Fig. 4C and Supplementary Fig S3**). Together these data indicate that early suppression of the IFN-system occurs below certain replication threshold, presumably mediated by virally encoded immune antagonists. This threshold is overrun by TMPRSS2-mediated surface entry, resulting in enhanced delivery of viral genomes to the cytosol to seed initial replication and concomitant early innate immune activation. Furthermore, these data suggest that the loss of tropism for A549 lung epithelial cells by B.1.1.529 is not due to an inability to antagonize the IFN system, as broad induction of antiviral responses, which supresses replication and abrogates virion release, was not observed. Indeed, at high MOI infection B.1.1.529 of A549-AT cells, replication overshoots this threshold and innate immunity is activated

### Initial entry route determines virus production kinetics and cell life-span

To investigate whether differences in initial viral uptake rates had downstream consequences on virion production and cell-death rates, we monitored particle production kinetics in time-course experiments. Distinct virion egress profiles were observed (**Fig. 4D**, left panel) which were likely facilitated by robust initial transfer of viral genomes to the cytosol, priming rapid innate immune activation in A549-AT cells, versus delayed induction in A549-A cells (**Fig. 4B** and **4C**). Virus release from A549-AT cells was initially higher but this pattern was reversed at later time points, with enhanced production observed from A549-A cells. Indeed, cumulative virus production was greater in A549-A cells (**Fig. 4D**, right panel). In line with this observation, A549-AT cell viability was reduced at 96 hpi when compared to A549-A cells (**Fig 4E**). These data point to a reduced life-span for highly permissive A549-AT cells, and highlight differences in the temporal dynamics of particle production and virus induced cell-eath which are modulated by initial entry route, determined by the presence or absence of surface TMPRSS2 expression.

### Innate immune activation status modulates the frequencies of specific mutations in NSP3

Upon transmission, fixation of mutations in RNA virus populations can occur through founder effect (limited genetic variability in the inoculum) or selective sweeps (an advantageous phenotype is swept to fixation) (For examples see (***Gömer et al., 2022***) and (***Brown et al., 2012***)). As the dynamics of early innate immune activation differs between viral entry routes, we hypothesized that the resulting contrasting selective environments, in which early viral replication occurs, could influence the frequency or fixation of mutations in the cellular viral population. To this end, we analyzed single nucleotide variant frequencies in cell-resident viral genomes and transcripts from our panel of infected cell-lines, focusing on non-synonymous substitutions. Generally, amino acid consensus sequences in all virally encoded ORFs remained highly conserved upon transmission. Non-synonymous substitutions with frequencies >3% were limited to a hand-full of residues distributed throughout the genome (**Fig 5A-C**). Interestingly, two residues (930 and 936) in close proximity in the primary amino acid sequence and located in ORF1a/ORF1ab exhibited shifts in the dominant amino acid. The observed shifts at these two positions appeared to be correlated to entry route, were reproducible between three biological replicates and occurred in all three strains in parallel (**Fig 5A-C**).

**Figure 5.**
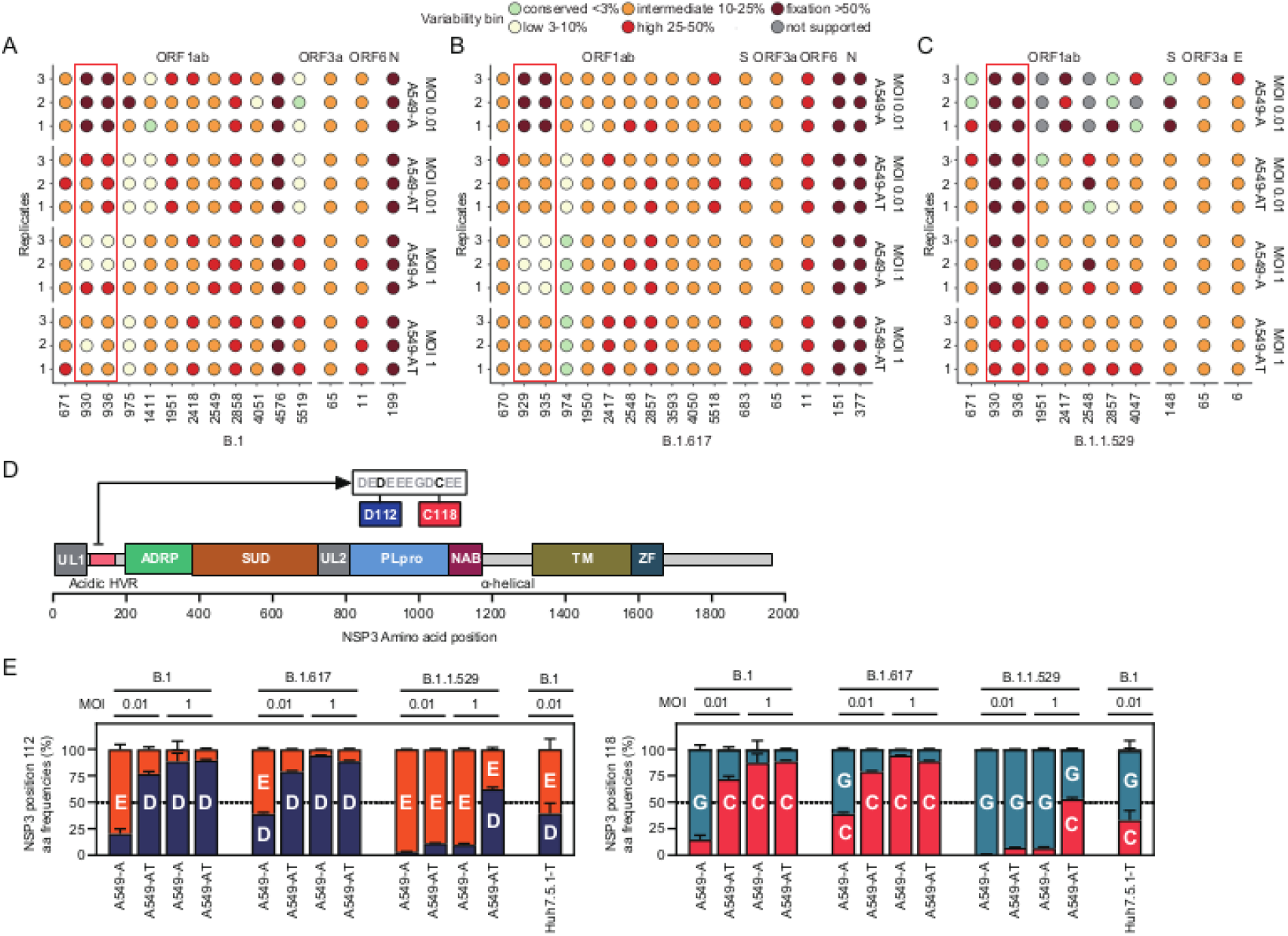
Viral NSP3 mutation frequencies at two residues are correlated to the innate immune status of infected cells. Dot plots highlight amino-acid variants above 3% frequency in the viral population, relative to the reference strain sequences for B.1 (**A**), B.1.617 (**B**) and B.1.1.529 (**C**). For each plot, biological replicates are displayed on the left y-axis and cell-line/MOI are presented on the right y-axis. The location of variable sites is displayed on the x-axis as amino acid numbering, with viral proteins in which they are located labeled above. For each position in the dot plot, the frequency of variability at that position is colored-coded relative to the key positioned above. Changes in the amino acid consensus (>50% frequency) are highlighted in dark red. Orf1a/1b residues 930/929 and 936/935 are highlighted by red boxes. (**D**) Cartoon depicting the protein domains of NSP3. UL: ubiquitin-like domain, HVR: hypervariable region, ADRP: ADP-ribose phosphatase, SUD: SARS-unique domain, PLpro: papain-like protease, NAB: nucleic acid binding domain, TM: multi-pass transmembrane domain, and ZF: zinc finger motif. Zoom of an 11 amino acid stretch of the NSP3 acid HVR highlighting positions D112 and C118 is positioned above. (**E**) Amino acid frequency plots highlight shifts in dominant residues for NSP3 positions 112 (left) and 118 (right) for the indicated viruses and cell-lines.

Orf1a/Orf1ab are synthesized as polyproteins which are post-translationally cleaved by host and viral proteases into multiple mature proteins with distinct biological functions. Mutations at Orf1a/Orf1ab positions 930 and 936 gave rise to D112E and C118G amino acid exchanges located in the acidic hypervariable region (HVR) of non-structural protein 3 (NSP3), a large multifunctional protein containing multiple distinct domains (***Armstrong et al., 2021*; *Lei et al., 2018***) (**Fig. 5D**). In engineered A549 cells, D112E and C118G mutations only became dominant in the viral population when host antiviral responses were minimally activated. To confirm this, we also performed B.1 infections of Huh-7.5.1 cells ectopically expressing human TMPRSS2 (Huh-7.5.1-T) as these cells possess ablated antiviral responses. In this background of minimal innate immune induction, D112E and C118G mutations also predominated (**Fig. 5E**). Mutation frequencies at these two positions were linked, implying they are compensatory (**Figs. 5E**) and are correlated to the magnitude of the host response (e.g. see antiviral TFs induction in **Fig. 4B**). In our experimental system, this pair of viral mutations act as molecular sensor, conveying information about the innate immune activation status of infected cells. Strikingly, this property was conserved in ancestral and VOCs, pointing to evolutionary conservation of the viral acidic HVR due to a shared function between strains and possibly reflecting a direct binding of this region to unknown host factors.

### Evolutionary conservation of the TMPRSS2 transmembrane domain and serine protease catalytic triad

While our data point to an evolutionary conserved function of the virus NSP3 acidic HVR between strains, cellular factors necessary for entry also exhibit evolutionary conservation at the species level. SARS-CoV-2 represents a zoonotic pathogen of probable bat origin (***Zhou et al., 2020***), with evolutionary conservation of critical ACE2 binding site residues determining the species range (***Liu et al., 2021*; *Yan et al., 2021***). However, whether diverse species exhibit evolutionary conservation of critical TMPRSS2 domains essential for S2’ cleavage and enhancement of viral entry remains unknown. Therefore, to explore the broader relevance of our findings, we next investigated the species-breadth of TMPRSS2-mediated infection enhancement from selected species incorporating established SARS-CoV-2 animal models, potential zoonotic reservoir species and human companion animals. Is infection enhancement a unique property of human TMPRSS2 or can orthologues from other species also enhance SARS-CoV-2 entry to similar degrees? Answering this question may provide insights into the origins of the pandemic, highlight highly permissive nonhuman reservoir species and determine the suitability of animals models for recapitulating human pathogenesis phenotypes.

Firstly, we performed phylogenetic analyses of selected TMPRSS2 ORFs from diverse mammals and rooted with zebrafish. These data revealed evolutionary relationships based on TMPRSS2 sequences were broadly consistent with reported groupings of eutherian mammals (***Meredith et al., 2011***), although rodents were antecedent to the main placental grouping (**Fig. 6A**). Both membrane localization and protease activity are essential for TMPRSS2 entry enhancement. To confirm broad conservation of these features, bioinformatics analyses of TMPRSS2 orthologues from selected nonhuman species were performed. Sequence comparison of translated proteins confirmed complete conservation of serine protease HDS catalytic triad residues (**Fig. 6B**) while hydrophobicity plotting highlighted the presence of a single pass transmembrane domain in all species (**Fig. 6C**). These data confirm TMPRSS2 proteins from evolutionarily diverse species exhibit functional conservation of key domains necessary for viral spike cleavage.

**Fígure 6:**
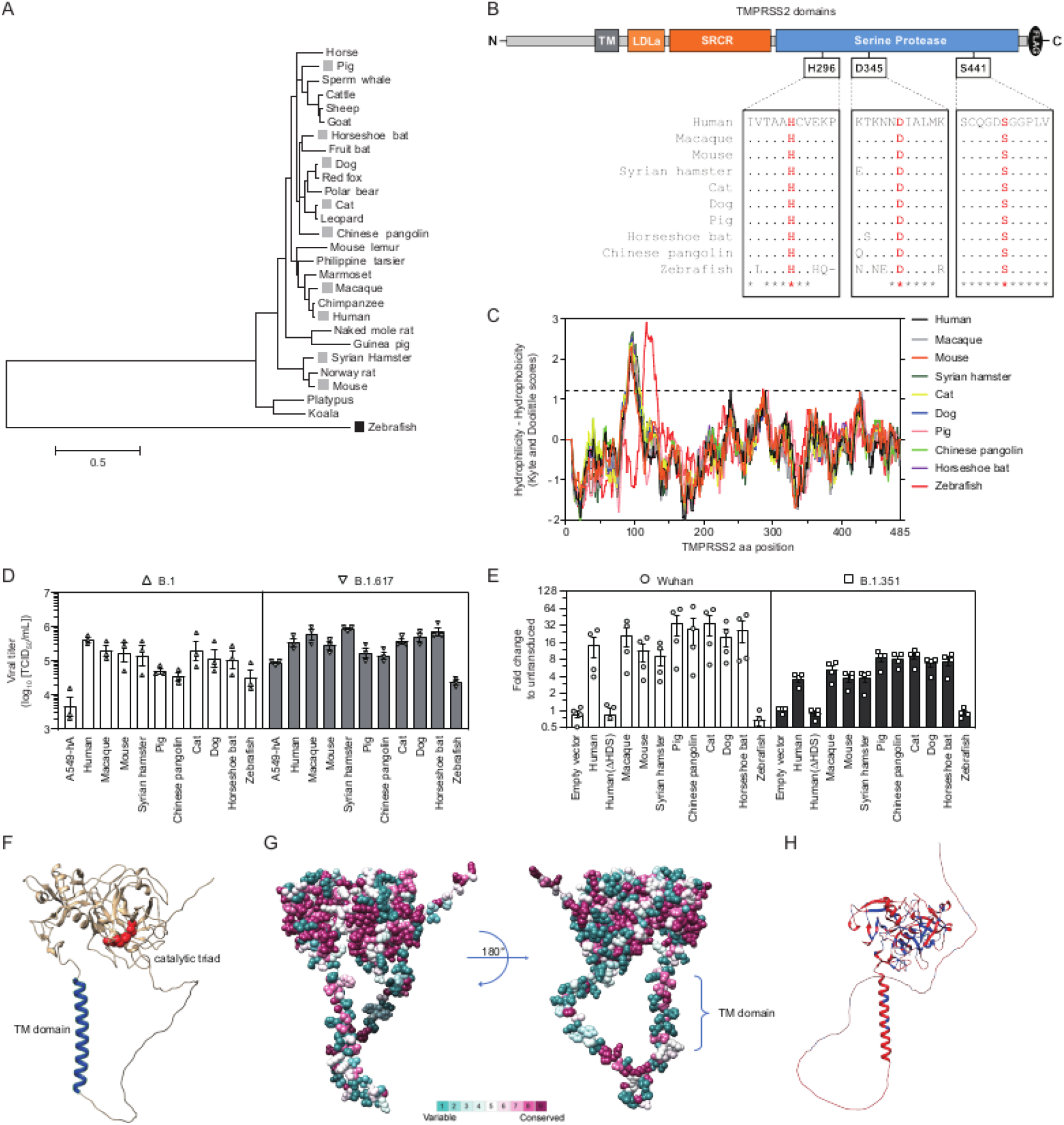
Evolutionary conservation of TMPRSS2 mediates entry enhancement in diverse mammals. **(A)** Phylogenetic tree depicts evolutionary relationships of *TMPRSS2* coding sequences. Branch length are proportional nucleotide substitutions per site, as determined by the scale bar. Species selected for further experimental investigation are highlighted. (**B**) Conservation of serine protease catalytic triad residues in humans and selected nonhuman species. (**C**) Hydrophobicity plots identify a single putative transmembrane domain in humans and nonhuman species. (**D**) Viron production rates at 24 hpi in A549-A cells co-expressing the indicated human and nonhuman TPMRSS2 orthologues. (**E**) SARS-CoV-2 pseudotyped vector transduction rates in HEK-293-ACE2 cells expressing indicated human and nonhuman TPMRSS2 orthologues. (**F-H**) TMPRSS2 structural predictions. (**F**) Human TMPRSS2 prediction with TM domain (blue) and HDS catalytic triad residues (red) highlighted. (**G**) Amino acid variability in mammalian TMPRSS2 orthologues superimposed onto the human TMPRSS2 structural prediction. (**H**) Amino acid differences (red) and conservation (blue) between mammalian and zebrafish TMPRSS2 orthologues are superimposed onto the zebrafish TMPRSS2 structural prediction.

### TMPRSS2 orthologues from diverse mammals enhance SARS-CoV-2 entry

Infections of A549-A cells ectopically expressing nonhuman TMPRSS2 orthologues were performed and virion production rates at 24 hpi compared to control A549-A and A549-AT cells (**Fig. 6D**). For B.1 infections, all mammalian TMPRSS2 orthologues enhanced virus production, ranging between 1-2 logs. This trend was less pronounced for B.1.617, with minimal enhancement observed for some species. Evolutionarily most distant, zebrafish TMPRSS2 was the least efficient at enhancing virion production for both strains. However, due to rapid SARS-CoV-2 replication kinetics, assessment of direct entry effects mediated by TMPRSS2 may be blurred by secondary rounds of infection and other confounding factors including suppression by the IFN system. Consequently, to complement the authentic virus infection data, HEK-293T-hACE2 cells ectopically expressing nonhuman TMPRSS2 orthologues were transduced with lentiviral pseudotyped vectors decorated with SARS-CoV-2 spike proteins from ancestral and VOC strains (**Fig. 6E**). Robust 4- to 20-fold entry enhancement for all mammalian orthologues was observed, while no enhancement was observed for the ΔHDS mutant or zebrafish TMPRSS2. Furthermore, for both authentic virus infections and pseudotyped vector transductions, ancestral strains appeared more dependent on TMPRSS2 for efficient entry than more recent VOCs (**Fig. 6D** and **6E**).

Despite transmembrane domain and catalytic triad conservation, zebrafish TMPRSS2-mediated entry enhancement was either highly inefficient (**Fig. 6D**) or non-existent (**Fig. 6E**). To investigate this in more detail, we visualized artificial intelligence (AI) generated structural predictions of TMPRSS2 proteins encoded by the human and zebrafish genomes (***Jumper et al., 2021***). Predicted structures display obvious TM helices while discontinuous HDS residues reside in close proximity in the globular serine protease domain (**Fig. 6F**). Residue variability in primary amino acid sequences for diverse mammalian orthologues was mapped onto the human structure, highlighting conservation of the serine protease domain located on the cell surface (**Fig. 6G**). The predicted zebrafish TMPRSS2 structure was used as a scaffold to visualize differences in primary amino acid sequences that are conserved between mammals and distinct in zebrafish (**Fig. 6H**). These analyses highlight differences which contribute to TMPRSS2 infection enhancement by comparing enhancing and non-enhancing orthologues in a structural context, which appear to be independent of transmembrane localization or the presence of a catalytic triad.

### *Tmprss2* gene transcription or mRNA abundance in hamster lungs is significantly down-regulated upon SARS-CoV-2 infection

SARS-CoV-2 infections of engineered A549 cells demonstrate viral entry route impacts a broad range of downstream host and viral processes *in vitro* in a homogenous lung epithelial cell population. Infections *in vivo* are seeded in the pharynx or lung, complex tissues composed of a heterogeneous mix of cell-types. The Syrian hamster has become the gold-standard animal model for SARS-CoV-2 infection research as it recapitulates many disease phenotypes seen in humans. In agreement with this, we demonstrate conservation of hamster TMPRSS2-mediated entry enhancement of SARS-CoV-2 (**Fig 6**). Consequently, to investigate parallels or differences between our *in vitro* RNA-seq analyses of infected A549 cells and *in vivo* infections of lung tissue, we leveraged bulk and single-cell (sc) RNA-seq data sets from our recent hamster vaccination studies (***Ebenig et al., 2022*; *Hörner et al., 2020***). Endogenous transcript levels in uninfected hamster lungs revealed baseline expression levels of infection-relevant genes *in vivo* (**Fig. 7A**). We observed comparatively low expression of *Ace2* transcripts, while pro-entry factors *Tmprss2*, *Ctsv* (homologous to *CTSL* in humans) and *Ifitm2* were abundantly expressed. A recent report describes Fcγ receptor mediated uptake of IgG opsonized SARS-CoV-2 as an alternative route of virus uptake into monocytes (***Junqueira et al., 2022***) and we noted intermediate expression of *Fcgr2b* and *Fcgrt.* Low-to-abundant expression of multiple upstream regulators of the host antiviral response was also apparent (**Fig. 7A**). As previously, infection-induced DEGs were identified by comparing uninfected and B.1 infected lung transcriptomes. Of note, further down-regulation of *Ace2* and significant down-regulation of *Tmprss2* were observed upon B.1 infection (**Fig. 7B**). This pattern may not represent gene regulation *per se*, but may rather reflect infection-induced death of permissive (*Ace2^+^*) and highly permissive (*Ace2+Tmprss2+*) cells, resulting in depletion of these transcript levels in bulk sampled tissue (as demonstrated experimentally in Fig. 4). In contrast, expression of transcripts encoding entry promoting factors *Ctsv* and *Ifitm2* and Fcγ receptors *Fcgr2b and Fcgr4* were significantly upregulated upon infection. Similar to engineered A549 cells, significant upregulation of transcripts encoding chemokines, interleukins, antiviral TFs and effectors was observed. In contrast to A549 cells, significant dysregulation of acute phase response TFs, nuclear receptor TFs and type I and III IFNs was not observed, likely reflecting the later sampling point (4 dpi). In infected hamster lungs at 4 dpi, ~1% of reads mapped to the B.1 genome scaffold (**Fig. 7C**). Coverage mapping highlighted differences in viral transcription rates apparent between engineered A549 cells and infected hamster lungs (**Fig. 7D**). While tiled transcript profiles are similar in the 5’ portion of the genome encoding the non-structural proteins, a noticeable drop-off in hamster lungs was observed in the 3’ portion encoding the structural proteins. This may point to differences in viral transcription kinetics between cell-lines and lung tissue, or may reflect directed suppression of structural gene transcription in lung tissue that is not recapitulated *in vitro*, or transcript degradation over time. Consistent with our hypothesis that acidic HVR residue frequencies relay information about innate immune activation status, D112 and C118 residues were dominant in lung resident viral RNAs and no evidence for increased frequencies of E112 or G118 were observed (**Supplementary Fig. S4**).

**Figure 7.**
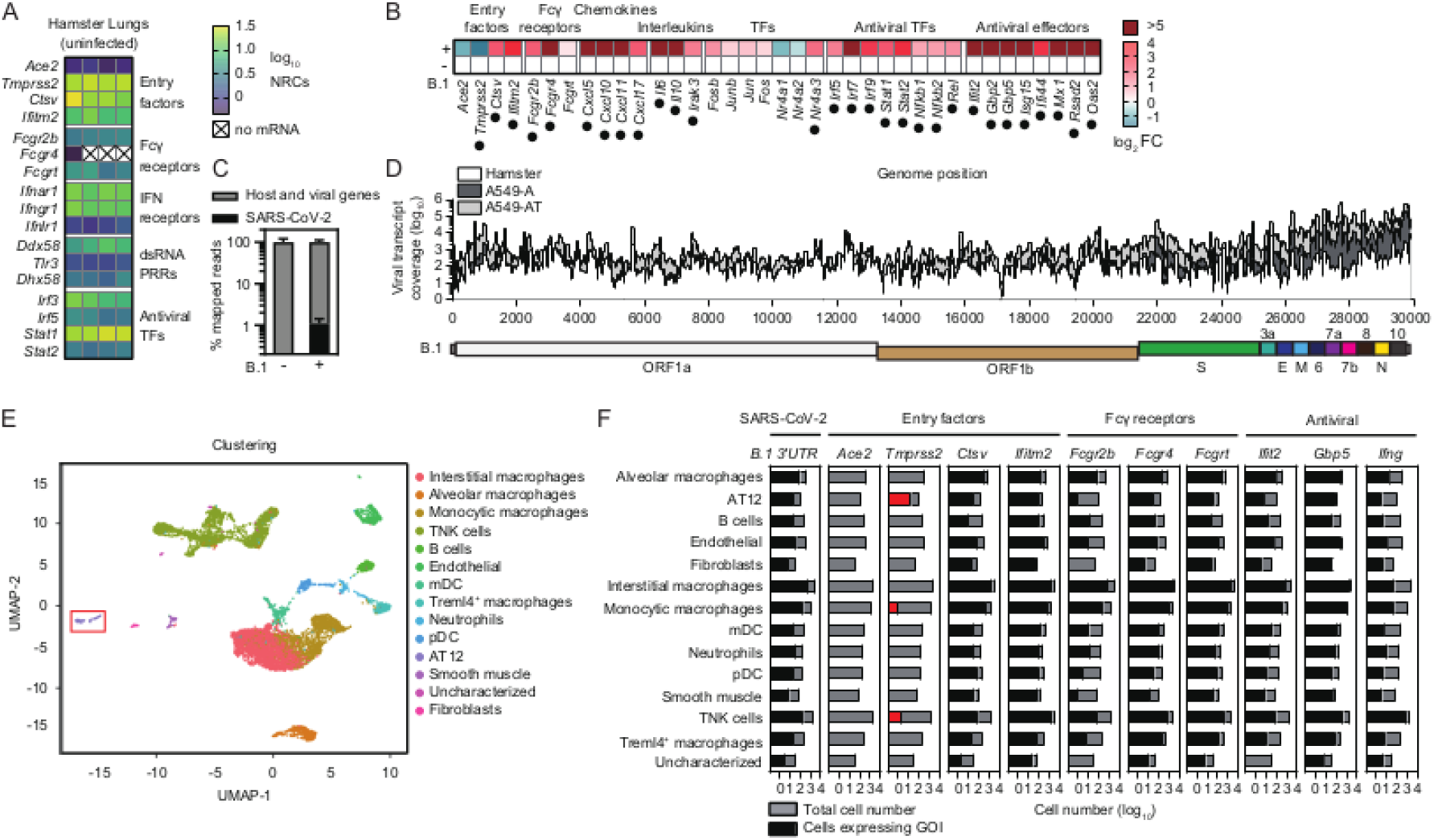
Hamster lung *Tmprss2* transcripts are depleted upon SARS-CoV-2 infection. (**A**) Heat map visualizes intrinsic expression of selected transcripts in the lungs of n=4 uninfected hamsters, determined by RNA-seq. NRCs. Normalized reads counts. (**B**) Heat map visualizes mean fold-change in lung resident mRNA expression (log_2_) for selected transcripts after SARS-CoV-2 infection of hamsters (n=4), relative to expression levels in uninfected control hamsters (n=4). (**C**) Number of virus-genome mapped reads as a percentage of total reads. (**D**) Mean genome/transcript coverage across B.1 genomes in A549-A and A549-AT cells (n=3), compared to profile in hamster lung (n=4). Cartoon directly below represents the relative locations of genome encoded ORFs. (**E**) UMAP plot reveals lung resident cell types. AT12 cells are boxed. (**F**) Cell-type expression of selected genes of interest (GOI).

### *Tmprss2* expression in hamster lungs is largely confined to pulmonary alveolar type I and II cells

To pinpoint the cellular contributors of infection-relevant genes in the lung, and identify cell types harbouring viral RNAs, we re-analysed scRNA-seq data from B.1 infected hamsters (***Ebenig et al., 2022***), identifying 13 distinct cell types (**Fig. 7E**). In agreement with low-level expression seen in bulk RNA-seq data, no *Ace2* expression was detected in any cell type (**Fig. 7F** and **Supplementary Fig. S5**). This may reflect down-regulation in infected cells or infection-induced cell-death combined with a requirement for deeper sampling to detect low abundance transcripts. *Tmprss2* transcripts were robustly detected only in subsets of pulmonary alveolar type I and II cells (AT12), consistent with previous reports in hamsters (***Nouailles et al., 2021***) and humans (***Muus et al., 2021***). Entry factors *Ctsv* and *Ifitm2*, and antiviral effectors (e.g. *Ifit2, Gbp5*) were broadly expressed across most cell-types, as were SARS-CoV-2 RNAs. *Fcgr2b, Fcgr4* and *Fcgrt* expression was enriched in macrophage populations, highlighting the potential for Fcγ mediated uptake as an alternative entry route in these cell-types. Of note, in contrast to A549 cells where type I and III IFNs were expressed, we detected broad cellular expression of type II IFN *Ifng*, which was enriched in the TNK cell compartment. Together these data highlight *Tmprss2*-mediated enhancement of infection *in vivo* would mostly be confined to AT12 cells in the lung, and highlights the range of cell types susceptible to infection despite low *Ace2* expression.

## Discussion

In our *in vitro* model, all tested SARS-CoV-2 strains exhibited some degree of TMPRSS2-dependency. We observed B.1.617 entry was less dependent on TMPRSS2, when compared to B.1 and B.1.1.529. These data contrast with reports showing that B.1.1.529 cell-entry has reduced TMPRSS2 dependency compared to B.1.617 (***Meng et al., 2022*; *Zhao et al., 2022***). In our hands, B.1.1.529 demonstrated reduced entry and lower replication in VeroE6 cells and engineered A549 cells, although enhancement of intracellular genomes/transcripts was observed in the presence of TMPRSS2 at both low and high MOI. Despite robust detection of cell resident genomes and transcripts in engineered A549 cells, B.1.1.529 egress was completely abolished: This altered cellular tropism partially masks the TMPRSS2 dependency of B.1.1.529. Together these data demonstrate a range of TMPRSS2 dependencies for ancestral and VOC strains, and a loss of lung epithelial cell tropism for VOC B.1.1.529 which appears to be independent of ACE2 receptor expression. These data point to either an absence of unknown dependency factors or the presence of potent restriction factors in this cell-type, which uniquely target the B.1.1.529 lifecycle. In our model, B.1.1.529 retains TMPRSS2 dependence upon entry, determined by enhanced nucleocapsid gene and protein expression (**Fig. 1**) and high-resolution transcriptional profiling (**Figs. 3** and **4**). In contrast to previous strains, B.1.1.529 also utilized murine ACE2 for entry, expanding the potential species range to rodents but also allowing for infection of standard laboratory mice.

In A549-A cells, only entry via endosomes is possible due to the lack of endogenous TMPRSS2 expression. In contrast, in A549-AT cells, surface or endosomal entry are both technically possible and we cannot exclude simultaneous entry by both pathways (***Ou et al., 2021***). However, surface entry is much more efficient than endosomal entry **(*Koch et al., 2021*)** and our data indicate at low MOI, most virions likely enter A549-AT cells via the surface route when receptor:protease complexes are not saturated. Indeed, infection of A549-AT cells was highly efficient and resulted in a short burst of intense viral propagation, concomitant with robust activation of innate immunity. At high MOIs, saturation of receptor: protease complexes may lead to virion engagement of solitary uncoupled ACE2 receptors, facilitating internalization to endosomes and simultaneous entry via both routes. At high MOIs, differences in viral replication/transcription rates and host responses between A549-A and A549-AT cells were minimal, likely due to saturation of available replication sites or co-opted factors within the cell. Under these conditions, entry dependent differences between the two cell-lines were masked (**Fig. 3**).

Our data confirm endolysomal entry is less efficient and demonstrates a concomitant decrease in early viral replication, transcription and virion release. However, at these levels, the virus is able to partially counteract cellular antiviral responses, and innate immunity is not robustly activated. These data confirm that SARS-CoV-2 can spread under the radar of innate immunity below a certain replication threshold. Extrapolating these *in vitro* data to real-life infections, we propose a model whereby highly permissive *ACE2^+^TMPRSS2^+^* cells are initially infected and become a reservoir that support rapid viral spread to less permissive *ACE2^+^* cells, which promote intermediate levels of viral propagation. While accelerated virus-induced cell death occurs in highly permissive cells, delayed cellular intrinsic immunity inadvertently leads to an extended period of viral shedding from less permissive *ACE2^+^* cells. Schuler et al. identified that *TMPRSS2* expression was highest in alveolar epithelial type I cells (AT1) and ciliated cells and increased with aging in humans and mice (***Schuler et al., 2021***). Indeed, our hamster lung data identifies enriched *TMPRSS2* expression in AT12 cells and depletion of lung *TMPRSS2* transcripts after infection (**Fig. 7**). This is consistent with a study demonstrating foci of infection are confined to regions where ACE2 and TMPRSS2 proteins co-localize: bronchioles and alveoli **(*Tomris et al., 2022*)**. Of note, we detected very low *Ace2* mRNA expression in hamster lung, which was further depleted upon infection and below the detection threshold of scRNA-seq. Despite this, viral RNAs were detected in a range of cell types. This data implies virus entry can occur despite very low levels of endogenous receptor expression, and may also occur via receptor independent routes, including entry of IgG opsinized virions to FcγR expressing cells (***Ebenig et al., 2022*; *Junqueira et al., 2022***), which are not recapitulated in our A549 cell model. Together our *in vitro* data shed light on how COVID-19 may initiate and spread in the lung, with subtly distinct virus production and innate immune activation profiles associated with *ACE2^+^* and *ACE2^+^ TMPRSS2^+^* expressing cells.

A number of preclinical candidates targeting both TMPRSS2 and cathepsin are available **(*Chen et al., 2021*; *Hoffmann et al., 2020*; *Mahoney et al., 2021*; *Ou et al., 2021*; *Shapira et al., 2022*)**. In our cell model, E-64d was more potent than camostat mesylate, significantly inhibiting entry of B.1 and B.1.617 strains in both A549-A and A549-AT cells. Differential expression levels of *TMPRSS2* and *CTSL* specific cell types and drug responsiveness should be taken into account when evaluating these data. Indeed, in hamster lungs, *Tmprss2* expression was largely but not exclusively confined to pulmonary alveolar type 1 and 2 cells (AT12) while *Ctsv* cell-type expression was higher and more broad (**Fig. 7 and Supplementary Fig S7B**). Whether drug candidates can target *TMPRSS2^+^* cells and efficiently prevent viral entry and spread to *TMPRSS2^-^* cells *in vivo* is unclear. Concerning therapeutic targeting of host entry factors, our data support simultaneous targeting of both TMPRSS2 and cathepsins in a combination regimen, which may prove synergistic (***Padmanabhan et al., 2020***).

In addition to differential activation of cellular processes correlated to entry route, our data also recorded reproducible modulation of viral NSP3 HVR mutation frequencies, which were conserved between ancestral and VOC strains. We propose that these shifts in mutation frequencies represent the molecular footprints of direct targeting and subsequent escape from an unidentified host antiviral effector protein. In the absence of robust innate immune responses, residues D112 and C118 confer a fitness cost and are selectively deleterious: E112 and G118 are dominant in the population. In contrast, when robust immune activation occurs, D112 and C118 are selectively advantageous, facilitating escape from host targeting, and their population frequencies move towards fixation. The glutamic/aspartic acid (E/D) rich region in NSP3 is intrinsically disordered and performs a currently unknown function, although E/D rich proteins are reportedly involved in DNA/RNA mimicry, metal-ion binding, and protein:protein interactions (***Lei et al., 2018***). Ultimately, identification of host-determinant(s) mediating this switch in NSP3 HVR residues 112 and 118 upon immune activation may illuminate the biological function of this region and how it contributes to the viral lifecycle.

To investigate the authenticity of current SARS-CoV-2 animal models and provide insights into potential zoonotic reservoir species, we investigated the species breadth of TMPRSS2-mediated entry. Spike cleavage at S2’ was a conserved property of mammalian TMPRSS2 orthologues from diverse species, but not zebrafish, which has implications for virus circulation in nonhuman species. Comparative analysis of enhancing and non-enhancing orthologues in a structural context revealed spike cleavage determinants likely map to multiple functional domains, in addition to absolute the requirement for a functional serine protease and plasma membrane incorporation. Early pandemic strains exhibited greater TMPRSS2-dependence than more recent VOCs, indicating ancestral progenitor viruses were likely highly dependent on this protease for spread in the natural reservoir species, and implying directed evolution in humans away from TMPRSS2 dependence. Of note, in the pseudotyped vector transduction assay, TMPRSS2 enhancement was reproducibly higher for orthologues from *Laurasiatherian* mammals (carnivores, ungulates, bats) when compared to *Euarchontoglires* (primates, rodents). These data further support a bat origin for SARS-CoV-2 and imply intermediate host species may also reside in this superorder of placental mammals. Together, many mammal species have the potential to shape the continuous evolution of SARS-CoV-2, while the origins of the initial spillover to humans remain focused on the *Laurasiatheria* clade.

In summary, our data demonstrate differences in TMPRSS2 dependence between B.1 and VOCs B.1.617 and B.1.1.529 correlate with differences in downstream activation of innate immunity after viral entry, which are coupled to viral RNA levels in infected cells. TMPRSS2 therefore represents an important modulator of downstream cellular responses to infection and can determine the efficacy of inhibitors targeting cellular proteases or endosome acidification. The footprints of differentially activated host responses are recorded in viral genomes and manifest as switches in dominant amino acids at two NSP3 residues. These data also identify a previously unappreciated inverse relationship between increased efficiency of cellular entry mediated by TMPRSS2 and reduced duration of productive viremia mediated by enhanced antiviral responses and accelerated cell-death. We show that TMPRSS2-mediated entry enhancement is conserved in gene orthologues from diverse mammals, providing insights into zoonotic reservoirs and experimental models. This enhancement is increased for early pandemic strains when compared to VOCs. Aspects of our model are confirmed from *in vivo* infection data of hamsters. Finally, our data confirm both accessory proteases involved in SARS-CoV-2 entry represent viable therapeutic targets, and support combinatorial targeting of TMPRSS2 and cathepsins simultaneously to maximize potency and reduce spread in the lung.

## Materials and methods

### Generation of stable cell lines

*ACE2* and *TMPRSS2* orthologues used in this study were downloaded from Ensembl or GenBank and chemically synthesized (Integrated DNA Technologies). *ACE2* orthologues were ligated into the lentiviral pTsin-IRES-Puro vector via restriction digestion cloning. C-terminal FLAG-tagged *TMPRSS2* orthologues and the human ΔHDS mutant were ligated into the lentiviral pWPI-BLR vector (Addgene) via restriction digestion or using HiFi Builder (NEB). All inserts were validated by Sanger sequencing (Eurofins). Lentiviral plasmids were packaged into pseudoparticles via triple-plasmid co-transfection into HEK293T cells (TaKaRa). pTsin or pWPI plasmids, psPAX2 (HIV-1 gag/pol) (Addgene) and pMD2.G (encoding VSV-G) (Addgene) were transfected in equimolar amounts using Lipofectamine 3000 (Invitrogen). Supernatants were harvested at 24 and 48 hours after sodium butyrate induction (10mM, 6 h), pooled, filtered through 0.45 μM pores and stored at −80°C. Parental A549 cells were transduced with hACE2 and mACE2 encoding pseudotyped lentiviral vector particles. Puromycin (2 μg/mL; Sigma-Aldrich) was added to media preceding a two-week selection of A549-hACE2 (A549-A) and A549-mACE2 cells (A549-mA). A549-A cells were further transduced with pseudotyped lentiviral vector particles encoding TMPRSS2 orthologues or the ΔHDS mutant, and stabilized by blasticidin (20 μg/mL; Fisher Scientific) for two weeks, generating A549-A-hTMPRSS2 (A549-AT), A549-A-hTMPRSS2(ΔHDS) (A549-AT(ΔHDS)) and A549-A-xTMPRSS2 cells (x=macaque, mouse, Syrian hamster, pig, Chinese pangolin, cat, dog, horseshoe bat or zebrafish). Similarly, A549 and Huh7.5.1 cells were transduced and blasticidin selected to generate A549-hTMPRSS2 (A549-T) and Huh7.5.1-hTMPRSS2 (Huh7.5.1-T) stable cells.

### Cell lines

A549 and Huh7.5.1 cells were maintained in Dulbecco’s modified Eagle’s medium (DMEM) (Invitrogen) supplemented with Penicillin-Streptomycin, 1 mM L-glutamine (Invitrogen) and 10% fetal bovine serum (FBS) (Invitrogen). For maintenance of VeroE6 cells (ATCC CRL-1586), FBS was reduced to 5%. For expansion and culture of stable cell lines, the appropriate antibiotics were supplemented into the media, as described above. HEK-293T-hACE2 cells were grown in complete DMEM medium supplemented with zeocin (50 μg/mL; InvivoGen) (***Glowacka et al., 2010***).

### Viruses

Viral strains used in this study include the D614G B.1 isolate (MUC-IMB1, GenBank accession LR824570, a kind gift from the Institute for Microbiology, Bundeswehr), Delta B.1.617 (20A/452R) isolate (imported from Robert Koch Institute) and Omicron B.1.1.529 BA.1 isolate (FFM-SIM0550/2021, GenBank accession: OL800702) (***Wilhelm et al., 2022***). Virus stocks were propagated on VeroE6 cells and titers determined on VeroE6 cells using a TCID_50_ endpoint assay.

### SARS-CoV-2 infections

SARS-CoV-2 infections were conducted in 6-well or 12-well plates at MOI 0.01 or 1. Three to six biological replicates were performed for all infection experiments. Cells were inoculated with virus in serum-free DMEM medium for 1 h at 37°C with gentle shaking of plates every 15 mins. After infection, cells were washed 3x in PBS and complete DMEM was added. For inhibitors, cells were pre-treated with Camostat mesylate (10 μM; Sigma-Aldrich) or E-64d (10 μM; Santa Cruz Biotechnology) for 1 h prior to infection. After inoculation, these compounds were re-administered to the fresh medium. Bafilomycin A1 (1 μM; Santa Cruz Biotechnology) was used for 1 h pre-treatment but not for post treatment due to high cytotoxicity.

For virus titrations, 3×10^4^ VeroE6 cells per well were seeded in 96-well plates. The next day, a series of 1:10 dilutions of virus stocks or supernatant from infection experiments were prepared by serially diluting 30 μL of sample with 270 μL serum-free DMEM medium in 96-well plates, prior to transfer to seeded VeroE6 cells. Infected cells were incubated for 4 days at 37°C. Cytopathic effects in each well were identified using a phase contrast light microscope and documented. TCID_50_/mL values were computed using a TCID_50_ calculator.

### SARS-CoV-2 pseudotyped vector assay

Constructs expressing Wuhan and B.1.351 S proteins of were generated as previously reported (***Hastert et al., 2022***). SARS-CoV-2 pseudoparticles were produced by co-transfecting HEK293T cells using Lipofectamine 2000 (Invitrogen) with plasmids encoding HIV-1 gag/pol, rev, a luciferase-encoding lentiviral transfer vector, and the S gene from either SARS-CoV-2 Wuhan (614D; GenBank: MN908947.3) or B.1.351 (Beta variant). Pseudotyped vectors were harvested, concentrated by ultracentrifugation and stored at - 80°C prior to transduction.

HEK293T-hACE2 cells (3.0 × 10^3^/well) were seeded in 384-well plates and transduced with lentiviral vectors encoding xTMPRSS2 (x=human, human(ΔHDS), macaque, mouse, Syrian hamster, pig, Chinese pangolin, cat, dog, horseshoe bat or zebrafish) at an MOI of 1, normalized by RNA copy numbers in supernatant. At 48 h post transduction, cells were infected with SARS-CoV-2pps at volumes that resulted in 2 × 10^5^ relative light units in a previous titration. At 48 hpi, entry efficiency was determined using britelite plus luciferase substrate (Perkin Elmer), which was added to each well to quantify luciferase activity using a Tecan Spark luminescence reader (Tecan).

### Western blot analysis

Adherent cells were washed with pre-cooled PBS and lysed in modified radio-immunoprecipitation assay (RIPA) buffer (50mM Tris-HCl [pH = 8.0], 150mM sodium chloride, 0.1% SDS, 1% Nonidet P-40, 0.5% sodium deoxycholate) supplemented with protease inhibitor cocktail (Cell Signaling Technology) for 30 mins on ice. Lysates were transferred into Eppendorf tubes and centrifuged at 960×*g* for 10 mins at 4°C. Supernatant was preserved for western blot analysis. Protein content was determined by BCA protein assay (Thermo Scientific). Equal amounts for each sample were mixed with 4× protein loading buffer (200 mM Tris-HCl [pH = 6.8], 400 mM DTT, 8% SDS, 0.4% bromophenol blue, 40% glycerol), heated for 5 mins at 98°C, loaded onto a 4-20% pre-cast SDS gel (Bio-Rad) and resolved by SDS-PAGE. Proteins were subsequently blotted to a PVDF membrane, which was further blocked with 5% blotting-grade milk in TBS for 1 h at room temperature. The membrane was incubated with α-mouse ACE2 (cross-reactive to human ACE2) (1:1,000; #38241 Cell Signaling Technology), α-human TMPRSS2 (1:1,000; HPA035787 Sigma-Aldrich) or α-β-actin (1:1,000; Abcam) with a gentle agitation overnight at 4°C. The next day, washed membranes were incubated with horseradish peroxidase (HRP) coupled α-mouse and α-rabbit IgG(H+L) secondary antibodies (1:15,000, Jackson Laboratories). Bound antibodies were detected with ECL Plus Detection substrate (GE Healthcare) and visualized using a ChemiDoc Imaging System (Bio-Rad).

### Immunofluorescence

Cells were seeded in 24-well plates and infected with B.1, B.1.617 and B.1.1.529 at MOI of 0.01. At 24 hpi, cells were washed with PBS and fixed with 4% paraformaldehyde for 30 mins at room temperature. After three washes with PBS, cells were permeabilized for 30 mins in 0.25% Triton X-100 in PBS. After washes, cells were blocked with 5% milk in PBS for 1 h and then incubated with α-SARS-CoV-2 nucleocapsid antibody (1:3,000; #26369 Cell Signaling Technology) overnight at 4°C. After three washes, cells were incubated with Alexa Fluor 488 anti-rabbit IgG (H+L) (1:1,000; Life Technologies) and 4’,6’-diamidino-2-phenylindole (DAPI, 1 μg/mL) for 1 h at room temperature, protected from light. After washing, fluorescence signals were captured under a Nikon Eclipse Ti-S fluorescence microscope. Images were processed and merged using Fiji ImageJ software.

### RT-qPCR

Total cellular RNAs were isolated using a Direct-zol RNA Miniprep Plus Kit (Zymo Research) according to the manufacturer’s instructions. Total RNA concentrations were quantified using a NanoDrop 2000 spectrophotometer (Thermo Scientific). A total of 500 ng RNA was reverse transcribed into complementary DNA (cDNA) using a PrimeScript II 1st Strand cDNA Synthesis kit (TaKaRa) or a QuantiNova Reverse Transcript Kit (Qiagen) according to the manufacturer’s instructions. RT-qPCR of cDNA samples was performed using 2× Rotor-Gene SYBR Green PCR Mastermix (Qiagen) and Rotor-Gene Q real-time PCR cycler (Qiagen).

For absolute quantification of SARS-CoV-2 *N* gene copies, a 319 bp fragment containing partial N and RdRp genes was synthesized, digested with NheI and EcoRI and cloned into pcDNA3.1Zeo (ThermoFisher Scientific) to generate a pcDNA3.1Zeo-NRdRp plasmid for *in vitro* transcription. RNA amounts were quantified and 1×10^12^ copies of RNA were reverse transcribed. The cDNA product was 1:10 serially diluted and used as standards. For absolute quantification of human *ACE2* and *TMPRSS2* genes, pTsin-hACE2-IRES-puro and pWPI-hTMPRSS2-BLR plasmids were used as standards for each gene. For relative quantification of human *IFNB1* and *IFIT1* genes, calculations were performed using the ΔΔCt method and fold changes were calculated using the 2^-ΔΔCt^ method. For human genes, *GAPDH* was used for normalization. Validated primer pairs were taken from Primer3 (https://primer3.ut.ee/) or from (***Blume et al., 2021***).

### Acidification assay

Cells (1×10^6^) were pre-treated with concanamycin A (5 nM; Sigma-Aldrich) for 1 h at 37°C and inoculated with B.1 virus at an MOI of 0.01 in the presence of concanamycin A for 1 h at 4°C. The cells were washed with PBS and maintained with medium containing concanamycin A for 30 mins at 37°C. Subsequently, cells were incubated for 10 mins with pH = 5.0 or pH = 7.0 medium at 37°C. Fresh medium containing concanamycin A was given to the cells for an additional 3 h. Medium was changed and intracellular RNA and virus infectivity in the medium were measured after 24 h.

### Flow cytometry

2×10^6^ A549-A and A549-AT cells were infected with B.1 virus at an MOI of 0.01 for 96 h. Cells were trypsinized, pelleted and re-suspended in PBS. After washes with PBS, cells were fixed with 4% paraformaldehyde for 30 mins at room temperature, pelleted and re-suspended in FACS buffer (1% FBS in PBS). Cell re-suspensions were stained using the LIVE/DEAD Fixable Aqua Dead Cell Stain kit (1:1,000; Thermo Fisher Scientific) at room temperature for 30 mins. Stained cells were washed and analyzed by flow cytometry using the excitation wavelength 405 nM (LSRFortessa, BD). Raw data were analyzed using FlowJo v10 software.

### Phylogenetic analysis

Coding sequences from mammalian and zebrafish *TMPRSS2* orthologues were downloaded from Ensembl. Sequence alignment was performed in MEGA7 software (***Kumar et al., 2016***) followed by evolutionary analysis using the maximum likelihood composite approach implemented under the GTR+I+Γ model of nucleotide substitution. The significance of groupings was assessed using the bootstrap approach with 1,000 replicates. The phylogenetic tree with the highest log likelihood was presented with significant bootstrap values (>70%) assigned to the corresponding branches. The scale bar is proportional to the number of substitutions per site. The analysis incorporated a total of 28 sequences and 1,569 nucleotide positions were included in the final dataset.

### TMPRSS2 hydrophobicity analysis

Amino acid translations for 10 selected TMPRSS2 orthologues were pasted in the ProtScale tool in the Expasy (https://web.expasy.org/protscale). Hydrophobicity (Hphob) scores for each amino acid were calculated using the Kyte and Doolittle algorithm (***Kyte and Doolittle, 1982***).

### TMPRSS2 structure modeling

Coordinates for AI structural predictions of human and zebra fish TMPRSS2 proteins were acquired using AlphaFold (***Jumper et al., 2021***). Evolutionary conservation scores for individual amino acid residues in mammalian orthologues were projected onto the predicted human TMPRSS2 protein structure using the ConSurf server for protein alignments (***Landau et al., 2005***) using default settings. Highlighting of the catalytic triad residues and transmembrane domain onto human TMPRSS2, and mapping mammalian: zebrafish conservation onto zebra fish TMPRSS2 were performed using UCSF Chimera (***Pettersen et al., 2004***).

### RNA-Seq and scRNA-seq

Total cellular RNAs from were isolated using TRIzol reagent (Thermo Fisher Scientific) and RNA quality checking was performed using an Agilent Bioanalyzer. Polyadenylated RNA was isolated with the NEBNext Poly(A) mRNA module (NEB). We used a modified protocol of the NNSR priming method (***Levin et al., 2010***) to produce the cDNA libraries (***Brown et al., 2020***). Sequencing was performed on an Illumina NextSeq550 instrument with a 86 base, single-end setting, making use of a custom sequencing program, which omitted the detection steps in the first 2 sequencing cycles. Raw Fastq files were quality- and adapter-trimmed with *fastp* (***Chen et al., 2018***), mapping to the *hg38* human assembly, or to the corresponding viral genomes and gene-level read counting were performed with STAR (***Dobin et al., 2013***) using default settings. We used the DESeq2 *R* package (***Love et al., 2014***) for differentially expressed gene (DEG) analysis according to the following workflow. http://master.bioconductor.org/packages/release/workflows/vignettes/rnaseqGene/inst/doc/rnaseqGene.html). Gene set enrichment and KEGG pathways analyses were carried out with *clusterProfiler* (***Yu et al., 2012***) and *Pathview* (***Priel et al., 1990***) packages in *R*. Bulk and scRNA-seq data from B.1 infected hamsters was generated as previously described (***Ebenig et al., 2022***). Bulk RNA-seq data generated in this study were submitted to the NCBI GEO database and can be found under accession number GSEXXYYZZ.

### Viral diversity analysis

Trimmed reads were mapped against the reference sequences for B1, B.1.617 and B.1.1.529, respectively. An initial consensus sequence was created using the Sam2Consus tool to fill up N-stretches in references. The resulting consensus sequence was then used for downstream variant analysis. Briefly, the trimmed reads were mapped against the respective reference sequence using tanoti. The mapped files were then used as input to diversi-tools and vnvs-tools. Data visualization was performed with an in-house *R* script using the tidyverse library and ggpubr.

### Statistics

Student *t* and One-way ANOVA tests were performed in GraphPad Prism 9 on the data generated from biological experiments, with appropriate correction for multiple comparisons. *P<0.05, **P<0.01, ***P<0.001, ****P<0.0001 and no significance (n.s.) P>0.05.

## Acknowledgements

We thank Christian Pfaller for plasmids pTsin-hACE2-IRES-Puro and pTsin-IRES-Puro, Stefan Pöhlmann for HEK-293T-ACE2 cells, Eberhard Hildt and Innes Mhedhbi for the B.1.617 strain, and Jonathan K. Ball for spike plasmid B.1.351. RJPB, BSS and MDM are supported by BMG grant Charis 6a. MDM was further supported by the German Center for Infection Research (DZIF; TTU 01.805, TTU 01.922_00).

## Supplementary Figures

**Supplementary Figure S1.**
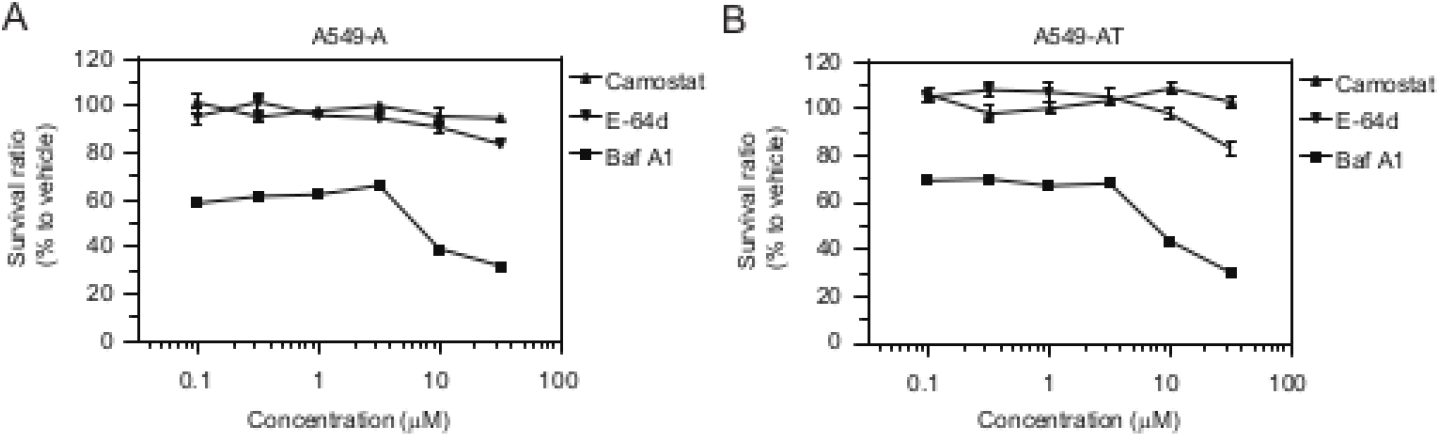
Cytotoxicity profiles of Camostat, E-64d and Bafilomycin A1 in A549-A (**A**) and A549-AT (**B**) cells.

**Supplementary Figure S2.**
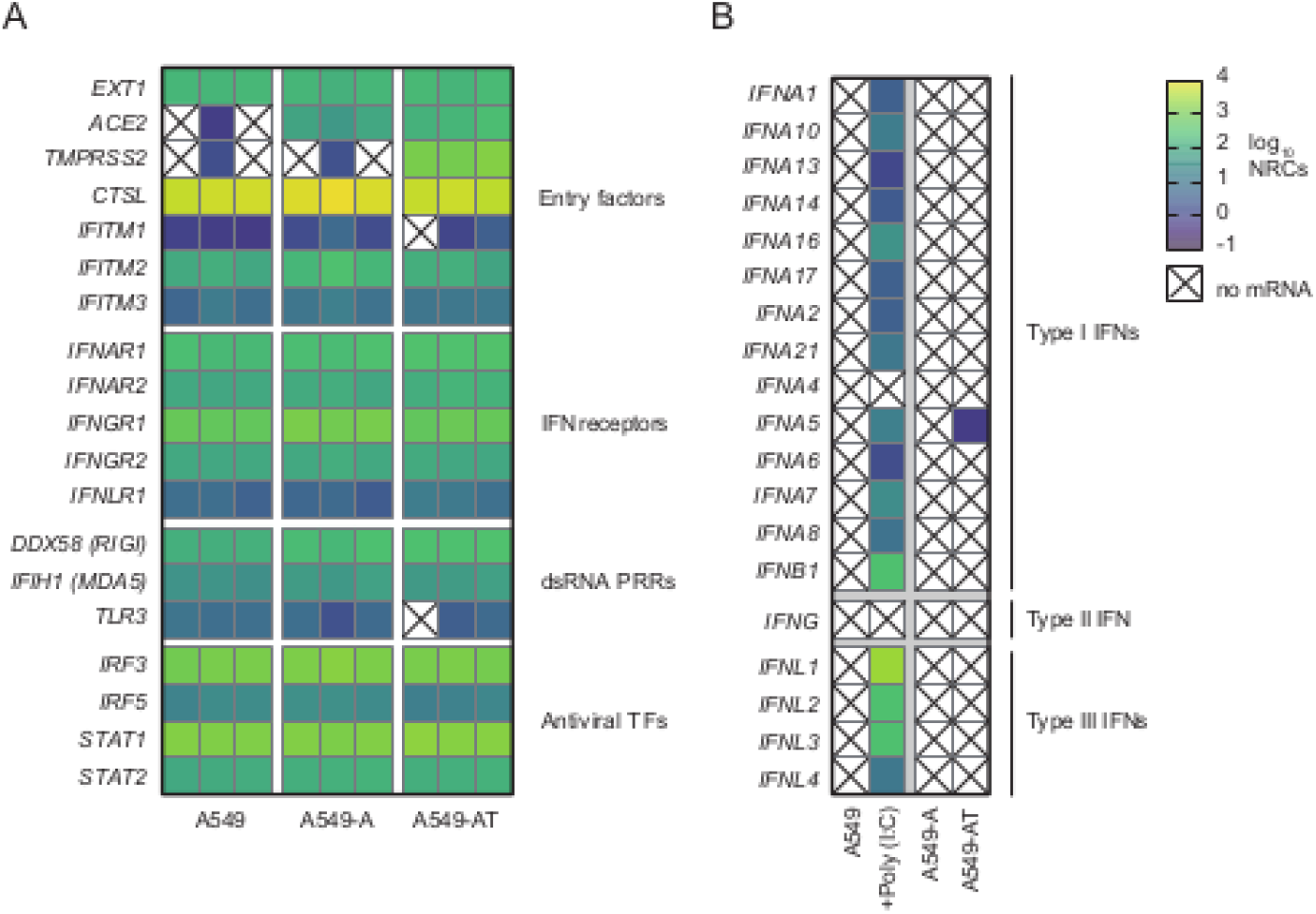
Intrinsic and inducible gene expression in parental and engineered A549 cells. (**A**) Transcriptional profiling of parental and engineered A549 cells without infection (n=3). Heat map visualizes intrinsic mRNA expression of viral entry factors, IFN receptors, dsRNA pattern recognition receptors (PRRs) and antiviral transcription factors (TFs) in the indicated cell-lines. Absent or minimal endogenous *ACE2* and *TMPRSS2* expression mRNAs is confirmed in parental A549 cells, while abundant ectopic expression is confirmed in engineered cells. (**B**) Robust and broad IFN induction after Poly(I:C) treatment of parental A549 cells (n=3, mean expression presented). Lentiviral gene transfer of *ACE2* and *TMPRSS2* does not induce IFNs. NRCs: normalized read counts.

**Supplementary Figure S3.**
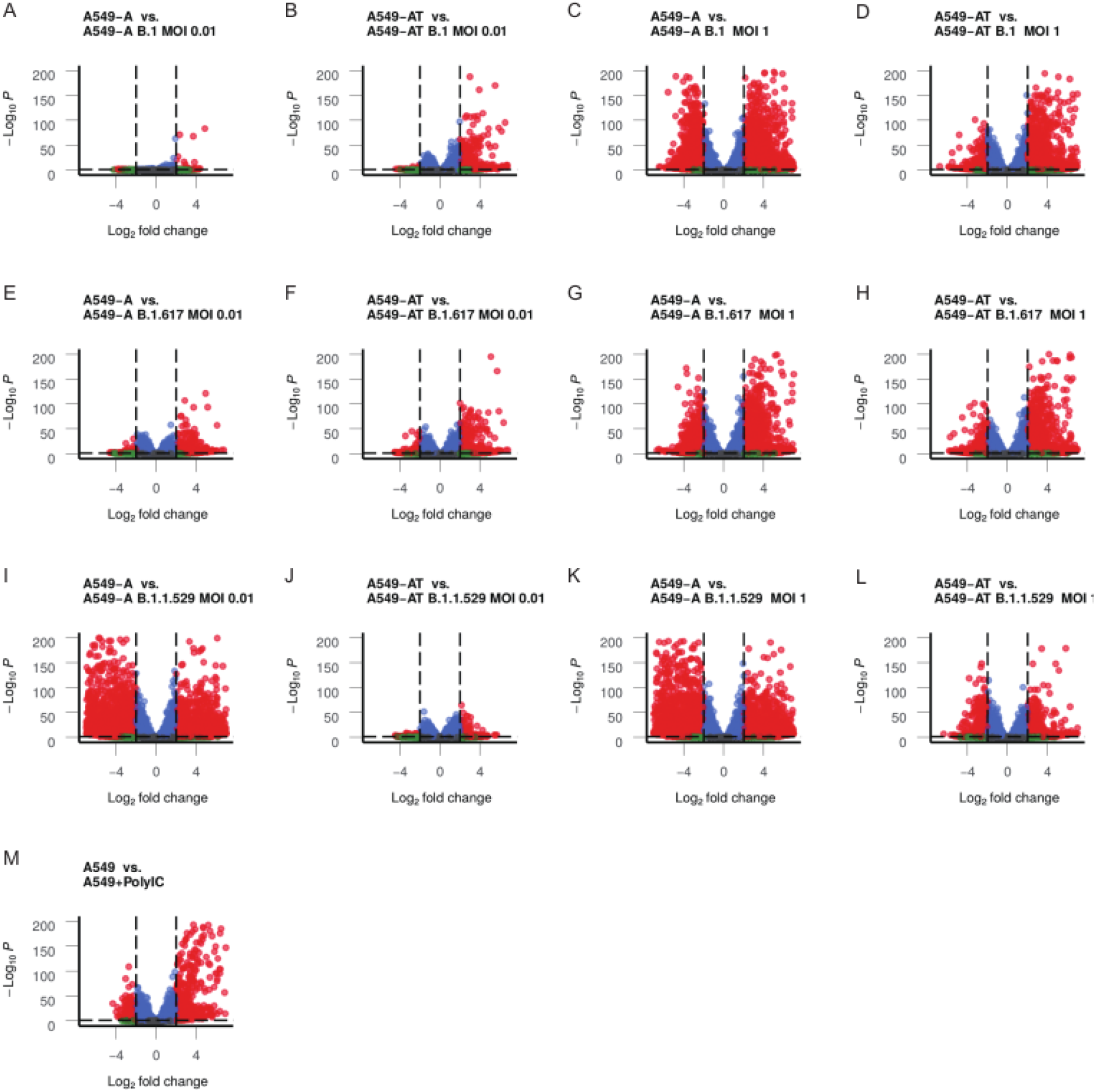
Volcano plots visualize differentially expressed genes induced upon SARS-CoV-2 infection or Poly(I:C) treatment of engineered or parental A549 cells, respectively. Infection of the indicted cell-lines with viruses B.1 (panels **A-D**), B.1.617 (panels **E-H**) and B.1.1.529 (panels **I-L**). Left two panels: MOI 0.01; right two panels: MOI 1. (**M**) Transfection of parental A549 cells with dsRNA mimic Poly(I:C). Thresholds for FDR p-values (y-axes) and log_2_-fold change (x-axes) were 0.05 and 2, respectively. Genes exceeding this threshold are highlighted red.

**Supplementary Figure S4.**
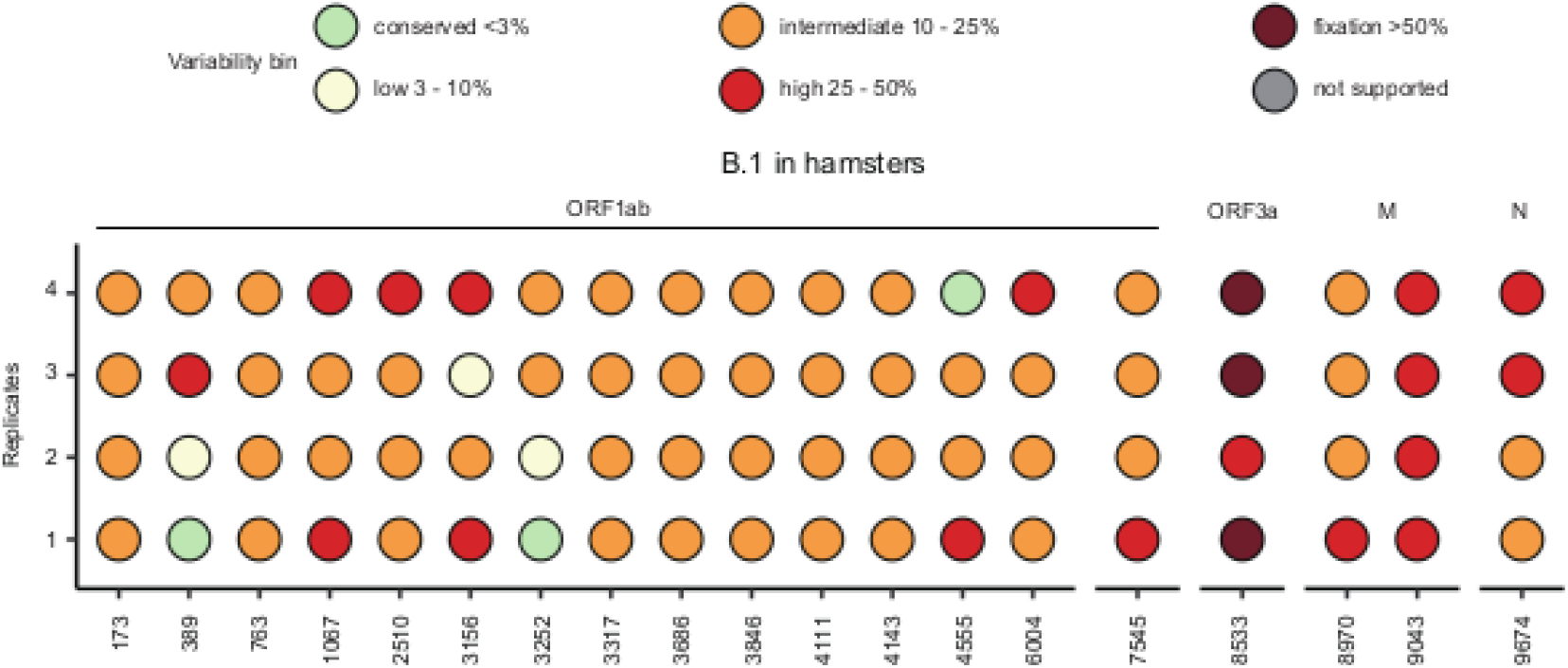
Dot plot highlights amino-acid variants above 3% frequency in hamster lung viral populations at 4 dpi, relative to the reference strain sequences for B.1. Biological replicates are displayed on the left y-axis and the location of variable sites is displayed on the x-axis as amino acid numbering, with viral proteins in which they are located labeled above. For each position in the dot plot, the frequency of variability at that position is colored-coded relative to the key positioned above. Changes in the amino acid consensus (>50% frequency) are highlighted in dark red.

**Supplementary Figure S5.**
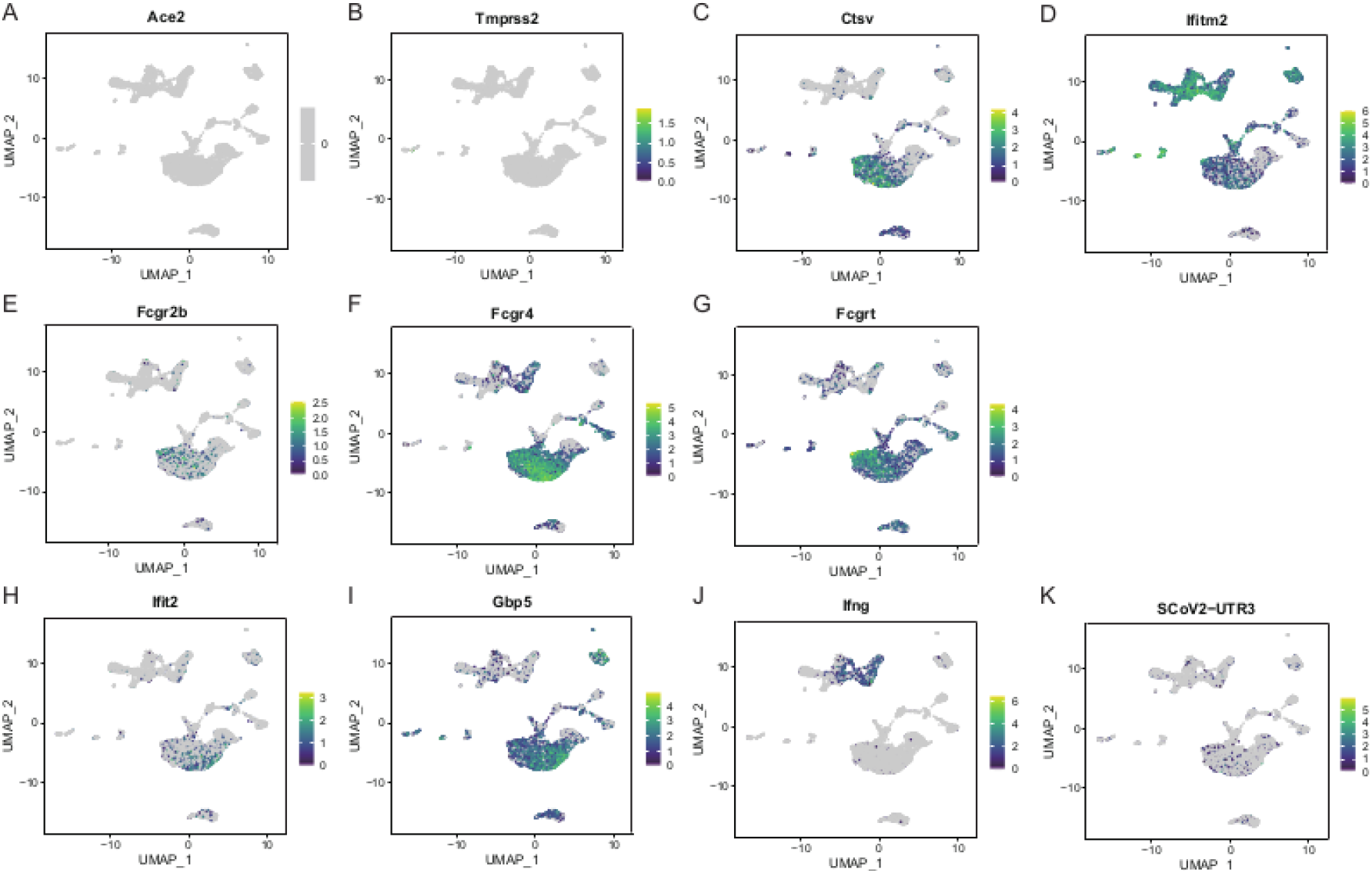
UMAP plot reveals lung cell-type expression of selected genes. Entry factor expression (**A-D**); FcγR expression (**E-G**); antiviral genes expression (**H-J**) and SARS-CoV-2 3’ UTR expression (K).

## Notes

### Competing Interest Statement

The authors have declared no competing interest.

